# Optical Spectral Fingerprinting Enables Sensitive Detection of Anthracycline Chemotherapeutics in Synthetic Clinical Biofluids

**DOI:** 10.64898/2026.04.08.717324

**Authors:** Atara R. Israel, Yunjung Kim, Adnan Arnaout, Myesha Thahsin, Yumna Ahmed, Zachary Cohen, Amelia Ryan, Syeda Rahman, Mijin Kim, Ryan M. Williams

## Abstract

Anthracycline chemotherapeutics are commonly used as frontline treatments for a wide array of cancers. However, their administration to patients results in substantial side effects, primarily cardiotoxicity, as well as myelosuppression and gastrointestinal toxicity. Current clinical management of such side effects is solely based on a lifetime dosage limit, which inhibits their anti-tumor efficacy. Many individualized factors, including age, family history of cardiovascular disease, treatment regimen, and other co-morbidities influence drug pharmacology. Despite this heterogeneity, there is no method for determining actual organ or tumor exposure to the treatment in an individual. Here, we developed an optical nanosensor array for four anthracyclines—doxorubicin, daunorubicin, epirubicin, and idarubicin. We used single-walled carbon nanotubes as the signal transducer due to their tunable near-infrared fluorescence. We screened twelve distinct ssDNA sequences paired with seven SWCNT *(n,m*) species at increasing concentrations of each of the four anthracyclines. The spectral responses were then used to develop machine learning-based classification models to identify different anthracycline types and concentrations. The optimized extreme gradient boosting model was able to classify high levels of each anthracycline with 100% accuracy. Concentration-based classification by PCA was performed for each anthracycline, distinguishing low (≤ 5 µM) and high (> 5 µM) concentrations. Finally, we validated the sensor performance using synthetic urine and sweat. Our findings demonstrate the potential of carbon nanotube-based sensor array to measure the pharmacokinetics of anthracyclines in patients with the goal of enhancing anti-tumor efficacy and monitoring off-target toxicities.

## Introduction

Anthracyclines are a potent class of chemotherapeutics derived from bacteria and widely used in the treatment of breast cancer, certain sarcomas, and leukemia/lymphomas^1-7^. Anthracyclines exhibit toxicity to both cancerous and healthy cells in several ways, including DNA intercalation, topoisomerase IIβ inhibition^8-10^, and cardiolipin antagonism^11^. Despite their efficacy, clinical anthracycline use is limited due to the lifetime dose-dependent cardiotoxicity and other side effects^8, 12-14^. These adverse effects can manifest both acutely and as long-term sequelae^15^. Current methods used to diagnose cardiac function include evaluation of left ventricle ejection fraction by echocardiography, or cardiac biopsy^16, 17^. Despite established guidelines^18^, anthracycline dosage is subject to substantial interpatient variability with a narrow therapeutic window. Thus, precise pharmacokinetic monitoring would be beneficial to optimize therapeutic outcomes while minimizing serious adverse events. Pre-clinical and clinical development methods for measuring anthracycline exposure in the body include mass-spectrometry^19^, high-performance liquid chromatography^20^, and the quantification of biomarkers of anthracycline-induced cardiotoxicity^16, 17^. These methods, however, are expensive, invasive, and/or may be too late for impact.

Semiconducting single-walled carbon nanotubes (SWCNTs) are quasi-one-dimensional nanomaterials comprised of sp^2^ hybridized carbon atoms. SWCNTs emit stable near-infrared (NIR) fluorescence,^21^ that is sensitive to changes in the local environment^22^. Many SWCNT chiralities exist, each defined by a distinct *(n,m)* index that specifies the nanotube diameter and chiral angle, each with individual absorbance and optical emission maxima^23^. The unique structure and electronic properties of *(n,m)* SWCNTs enable different interactions with their local environment.

Prior studies have used SWCNTs as transducers in sensors for a variety of clinically-relevant analytes, including small molecule neurotransmitters^24-28^, riboflavin^29-31^, and other chemotherapeutic agents such as cisplatin and transplatin^32^. SWCNT sensors have also been utilized to detect several biomarkers in patient samples, including estrogen receptor α^33^ and several gynecological markers^34, 35^. Implantable devices containing SWCNT have been engineered for in vivo sensor deployment to detect cancer markers^36^, steroid hormones^38^ and others^39^. Among chemotherapeutics, nanotube-based doxorubicin detection has been well investigated^37, 40, 41^. However, these SWCNTs function as generic anthracycline sensors rather than being specific to a single type^37, 40, 41^.

To improve the specificity of anthracycline sensing, we employed the spectral fingerprinting method^35, 42-44^. Spectral fingerprinting extends the molecular perception concept by correlating fluorescent responses from the sensor array with disease states, underlying cellular processes, or mixtures of analytes, and incorporates machine learning methods to discern these correlations. Here, we developed a SWCNT fluorescence spectral fingerprinting method to detect four structurally similar anthracyclines—doxorubicin, daunorubicin, epirubicin, and idarubicin (**Fig. 1**). The array consisted of twelve ssDNA sequences, some of which had been used for anthracycline detection and variants thereof, while we analyzed fluorescence intensity and wavelength changes from seven (*n,m*) species. We exposed the array to seven concentrations of each of the four anthracyclines in buffer, synthetic sweat, and synthetic urine. Fluorescence responses of ssDNA-SWCNTs were analyzed to differentiate among four anthracyclines, with the machine learning model achieving 100% classification accuracy. Quantitative sensitivity to each anthracycline, assessed using both traditional binding kinetic fits and machine learning models, achieved up to 100% accuracy in classifying concentration-dependent responses to daunorubicin and idarubicin. This work establishes the use of optical nanotube-based sensors that exhibit distinct responses to two anthracyclines, along with machine learning models to distinguish between different anthracyclines and to predict concentration.

**Figure 1.**
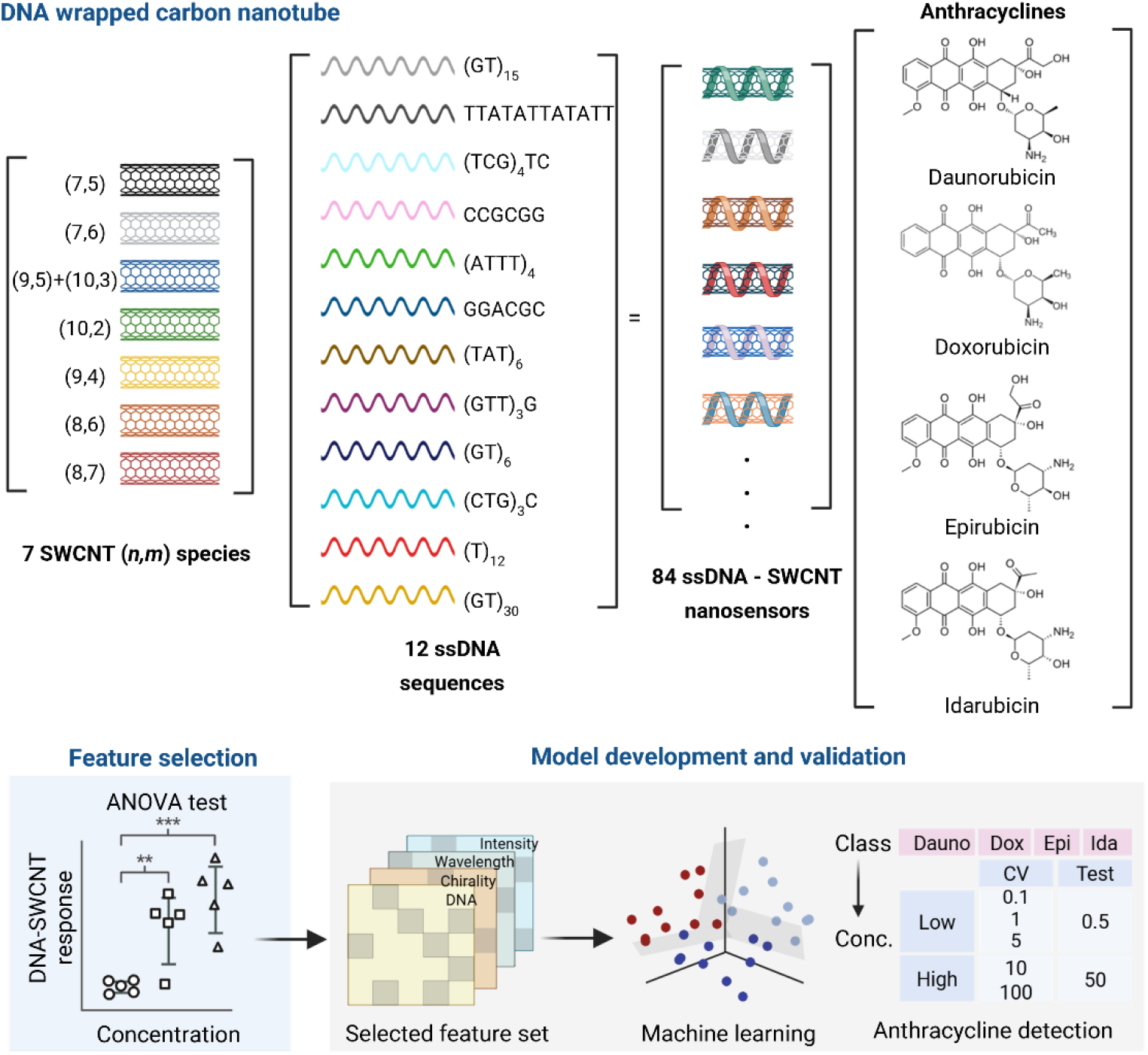
Workflow of nanosensor array screening and machine learning for anthracycline detection. Top: Dispersion of mixed-species SWCNT with ssDNA sequences, followed by addition of anthracyclines to each sensing platform. Bottom: Machine learning to determine optimal anthracycline sensors for classification based on anthracycline class and concentration.

## Methods

### Synthesis and characterization of the nanosensor array

As previously described^45^, twelve ssDNA sequences (**Table 1**; Integrated DNA Technologies; Coralville, IA) were combined, separately, in a 2:1 mass ratio with HiPCo SWCNT (NanoIntegris Technologies; Boisbriand, QC) in a final volume of 0.5 mL with 1X PBS (phosphate-buffered saline). The solution was then probe tip sonicated using a 3 mm stepped probe (Sonics Materials; Newton, CT) at 40% amplitude for one hour on ice. The mixture was then ultracentrifuged using an Optima Max-XP (Beckman Coulter; Brea, CA) at 58,000 X g for one hour. The top 75% of the supernatant was collected and stored at 4°C for future use. Prior to use, excess DNA was removed using a 100 kDa MWCO centrifugal filter (Sigma-Aldrich; St. Louis, MO) at 14,000 g for 15 minutes twice, with a washing step using 200 µL of 1X PBS between runs. The contents of the filter were resuspended in 1X PBS and characterized using absorbance spectroscopy over the range of 400-800 nm. The molar extinction coefficient at 630 nm of 0.036 L·mg^-1^·cm^-1^ was used to determine the SWCNT concentration of each suspension^46, 47^.

**Table 1.** List of ssDNA sequences and SWCNT (*n,m*) species analyzed in the engineered nanosensor array. Note: due to substantial overlap in the emission bands, the (9,5)+(10,3) species were analyzed as one.

| ssDNA sequences | ($n,m$) species |
| --- | --- |
| (GT)<sub>15</sub> | (7,5) |
| TTATATTATATT | (7,6) |
| (TCG)<sub>4</sub>TC | (9,5)+(10,3) |
| CCGCGG | (10,2) |
| (ATTT)<sub>4</sub> | (9,4) |
| GGACGC | (8,6) |
| (TAT)<sub>6</sub> | (8,7) |
| (GTT)<sub>3</sub>G |  |
| (GT)<sub>6</sub> |  |
| (CTG)<sub>3</sub>C |  |
| (T)<sub>12</sub> |  |
| (GT)<sub>30</sub> |  |

### Anthracycline addition to nanosensor array

The anthracyclines daunorubicin HCl (LC Laboratories; Woburn, MA), doxorubicin HCl (LC Laboratories), epirubicin HCl (LC Laboratories) and idarubicin HCl (Selleck Chemicals; Houston, TX) were solubilized in a 0.1% dimethyl sulfoxide (DMSO) solution. Additional heating was required to solubilize doxorubicin and epirubicin using a heating block at 60°C for eight minutes with 1000 RPM shaking. Each anthracycline was then added, separately, to 1 mg/L of each of the ssDNA-SWCNT suspensions in triplicate at concentrations of 100, 50, 10, 5, 1, 0.5, and 0.1 µM. These were incubated in a Corning UV-transparent half-area 96-well plate (Fisher Scientific; Hampton, NH) at a final volume of 150 µL in 0.1% DMSO for one hour at room temperature.

### High throughput NIR fluorescence spectroscopy

Fluorescence spectroscopy of the nanosensor array before and after anthracycline addition was obtained using a near-infrared plate reader (Photon, Etc.; Montreal, QC), with emission spectra acquired from 900-1600 nm after sequential laser excitation of 655 nm and 730 nm of 1700 mW and exposure time of 500 ms. Each laser excitation source was used to obtain spectra from (*n,m*) species: [(7,5), (7,6), and (9,5)+(10,3)], and [(10,2), (9,4), (8,6), and (8,7)], respectively.

### Data analysis

Changes in intensity and wavelength of SWCNTs (**Table 1**) were analyzed using a custom MATLAB code that fit each emission peak with a pseudo-Voigt profile to obtain the maximum intensity and center wavelength. Only R^2^ values of peak fits greater than 0.9 were used. Changes in center wavelength and maximum intensity were reported relative to the control (no anthracycline). The dissociation constants (K_d_) were derived from the spectral responses of SWCNTs using a standard non-cooperative binding model: 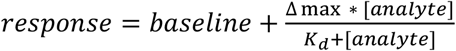 where response is the signal obtained (either change in center wavelength or intensity) at a given analyte concentration, baseline is the response of the control, Δmax is the maximum response observed, [analyte] is the analyte concentration, and K_d_ is the analyte concentration which elicits 50% of the maximum response. The fittings were performed in OriginPro. Statistical significance in fluorescence change was determined by a one-way ANOVA with Dunnett’s post hoc analysis using OriginPro. Principal component analysis (PCA) and machine learning models were implemented for classification in Python 3.12.2 using the *scikit-learn* 1.7.1 package^41, 48^.

### Multi-class classification of anthracycline type

Three supervised machine learning models, Decision Tree (DT), support vector machine (SVM), and eXtreme Gradient Boosting (XGBoost, XGB), implemented in the *scikit-learn* library, were used to develop multi-class classification models. The training set comprised spectral response data acquired at four concentration levels (0.1, 1, 10, and 100 µM). Model generalization and predictive accuracy were assessed using an independent test set comprising intermediate concentrations (0.5, 5, and 50 µM). Hyperparameter optimization was conducted using Bayesian optimization with *k*-fold cross-validation (*k* = 3) on the training set to identify the optimal model configuration. SHAP (SHapley Additive exPlanations) analysis was then applied to quantify the contribution of each spectral feature to the classification output^49^.

### Development and validation of concentration-based binary classification models

The training dataset was constructed exclusively from 1X PBS containing 0.1% DMSO at five anthracycline concentrations (0.1, 1, 5, 10, 100 μM). Test datasets were used as independent samples with intermediate concentrations (0.5, 50 μM). Concentrations ≤ 5 μM were labeled as “Low” and those > 5 μM as “High”. The model development and optimization process were the same as described in the multi-class classification models. Performance of the optimized models were assessed using 10% solutions of synthetic urine (Sigma-Aldrich; St. Louis, MO) and synthetic sweat (Sigma-Aldrich, St. Louis, MO) containing 1 mg/L of each sensor construct. The solutions were spiked with each anthracycline at concentrations of 1 and 50 µM. Sensor responses from each spiked medium were acquired and processed as described above. PCA was fitted on the training data only, and the resulting components were applied to both the training and test sets to avoid data leakage. These PCA-transformed features within the PC domain were used to discriminate between low- and high-concentration classes. Furthermore, SVM classification was performed on the PC domain to evaluate PCA-based discrimination accuracy. This validation quantified model robustness against matrix effects and compositional variability inherent in biofluids.

## Results and Discussion

### Construction and Characterization of SWCNT Sensor Array

SWCNTs were first encapsulated with 12 different ssDNA sequences, with subsequent analysis of 7 *(n,m)* structures to create 84 unique sensing elements (**Table 1**). ssDNA sequences were selected from the recognition sequences of SWCNTs for sorting and molecular sensing as previously reported. (GT)_15_ has been employed for anthracycline detection^37, 40^ and other small molecule detection^25, 30, 35, 50-54^, along with the shorter (GT)_6_ sequence^42, 50, 55, 56^. The (GT)_30_ sequence was included as a longer variation of these two poly-GT sequences and has been used in other SWCNT sensing applications^57^. Several of the sequences used in this study, including TTATATTATATT^58-60^, GGACGC^61^, (TCG)_4_TC^62^, (GTT)_3_G^62, 63^, (ATTT)_4_^64^ have been reported for use in chiral separation techniques, as certain nucleotide arrangements create unique motifs or conformations around the SWCNT surface. Variations of previously reported poly-CG sequences, CCGCGG and (CTG)_3_C were also included^61, 62, 65^. (TAT)_6_ was previously used for dispersion prior to antibody conjugation^33, 36, 47, 64, 66^ and is a variant of several poly-TA sequences for chirality sorting^62, 63^. (T)_12_ was included as it forms a somewhat weaker left-handed helical wrapping on SWCNT, allowing for more rearrangement^67^.

DNA-SWCNT complexes were characterized by UV-Vis absorption spectroscopy, followed by near-infrared (NIR) fluorescence spectroscopy using excitation wavelengths of 655 and 730 nm (**Supplementary Figure 1)**. Absorbance spectra demonstrated stable SWCNT suspensions. Some sequences, for example, (ATTT)_4_ and GGACGC, resulted in stronger dispersion and higher absorbance than others, such as CCGCGG. All DNA-SWCNT constructs produced bright emission peaks from several *(n,m)* species. Variance in brightness of *(n,m)* species demonstrated heterogeneity imparted by differential sequence interaction. As an example, (ATTT)_4_-SWCNT exhibited minimal fluorescence from the (8,3) and (9,5)+(10,3) species when excited at 655 nm. The (TAT)_6_-wrapped (7,6) SWCNT was dimmer, while the (8,7) species was brighter compared to other sequences evaluated. The (GT)_6_-wrapped (10,2) SWCNT were much brighter in comparison to all other ssDNA-SWCNT combinations. Predominant across all constructs evaluated, the (7,5), (7,6), and (9,4) species were the brightest at the excitation wavelengths used.

### Anthracyclines Induce Concentration-Dependent Spectral Responses of Nanosensor Array

SWCNT suspensions were exposed to doxorubicin, daunorubicin, epirubicin, and idarubicin at concentrations of 100, 50, 10, 5, 1, 0.5, and 0.1 µM (**Figure 2** and **Supplementary Figures 2-7**). We observed 20-40 nm redshifts for all ssDNA-SWCNT species assessed when incubated with 100 µM anthracycline, similar to our previous findings that studied the effect of doxorubicin on (GT)_15_-SWCNTs^37, 40^. Smaller magnitude redshifts, or slight blueshifts, were observed as the anthracycline concentration decreased. In contrast, intensity changes did not follow typical dose-response kinetics (**Figures 2A-D, Supplementary Figures 2-7**). While most (*n,m*) exhibited fluorescence quenching up to 80% in the presence of high anthracycline concentrations, (TAT)_6_-SWCNT brightness increased in response to daunorubicin and idarubicin across all *(n,m)* species. This result is similar to a previous report of a SWCNT doxorubicin sensor dispersed with the ssDNA sequence C_8_AGA_2_T_2_ACT_2_C_8_ (referred to as CB-13), which also observed brightening intensity and redshifting of up to 17 nm of the (7,6) species^41^. Other ssDNA-SWCNT combinations in that study, however, observed a trend of quenched intensity and up to 8 nm redshifts in response to doxorubicin. Our work indeed had overlapping sequence space with that study, notably (GT)_15_ and (ATTT)/(TAT) repeats, in which we found similar responses to doxorubicin. As that study did not investigate other anthracyclines, the divergence of responses for (TAT) but not (ATTT) repeats are notable. Interestingly, that study found a weak correlation between the ability of a sequence to suspend SWCNT and doxorubicin response^41^. Future studies may integrate both the library assembled here and the poly-C-flanked library from that work to enhance sequence space. In total, the divergence of responses is driven by the unique conformation each ssDNA sequence adopts in relation to the chiral angle and diameter of diverse *(n,m*) species^67, 68^. Uncoated SWCNT surface area allows direct analyte interaction^59^. Changes in fluorescence result from charge transfer and/or polarity modulation from the anthracycline^69-71^, with additional contributions from ssDNA restructuring^55, 67, 72^.

**Figure 2.**
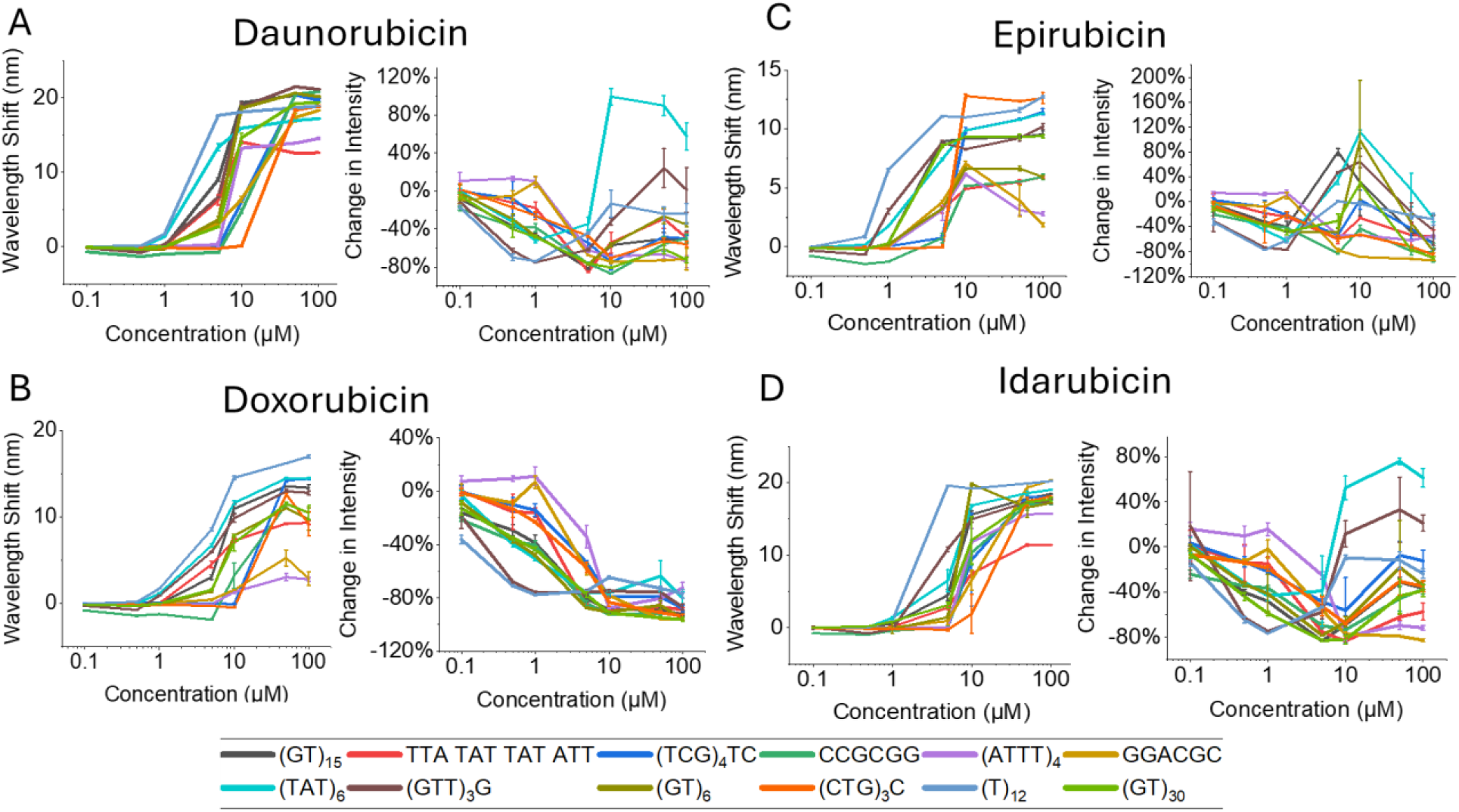
Concentration curves demonstrating shift in center wavelength and maximum intensity of (7,5)-SWCNTs. (**A**) daunorubicin, (**B**) doxorubicin, (**C**) epirubicin, (**D**) idarubicin.

Our prior work demonstrated that doxorubicin binding does not displace ssDNA on the nanotube surface^40^. The aromatic structure of anthracyclines, however, does have a strong affinity for the nanotube surface^73^. The affinity enables both direct charge transfer and polarity modulation. Anthracyclines are also strong DNA intercalators^74^, altering the ssDNA conformation on the nanotube surface and thereby fluorescence emission. This is supported by other studies which synthesized SWCNT-based electrochemical sensors with dsDNA, finding a reduction in voltametric peaks upon addition of anthracyclines, attributed to intercalation of the drugs with DNA^75-77^. Several other optically active nanomaterials, such as quantum dots and metal nanoclusters, have been designed for anthracycline detection, with the primary detection method being quenched absorbance or fluorescence. This is similar to our observations of SWCNT-anthracycline interactions here. The use of nanomaterials as a means to detect anthracyclines is a strong alternative to traditional methods due to the high affinity enabled by the surface chemistry of nanomaterials, and the lack of recognition elements for this drug class^78^.

We found the theoretical limits of detection (LOD; 3*σ_control_) ranged between 0.017-7.4 µM for wavelength-dependent modulations and between 0.1-5 µM for intensity-dependent modulations across ssDNA sequences and *(n,m)* species (**Supplementary Table 1**). The lowest wavelength-dependent LOD was obtained by the (TAT)_6_-(7,5), while the lowest intensity dependent LOD was obtained by the (TCG)_4_TC-(8,7). Interestingly, the highest LODs were observed in the (8,7) and (9,5)+(10,3) peaks, suggesting a trend of higher LOD for larger diameter species. Generally, we observed large wavelength shifts or intensity modulations in response to 10 – 100 µM anthracycline. At 5 µM, we observed notable variability, whereas some sensor combinations exhibited fluorescence modulations, others demonstrated almost no change. This is also true, to a lesser extent, at 1 µM. The addition of 0.5 and 0.1 µM anthracycline induced no substantial changes in fluorescence output. Other nanosensors for doxorubicin have found lower LOD, in some cases subnanomolar^78^. To cover a broad range of anthracycline concentrations with high accuracy, it is necessary to include multiple sensor constructs in a single model.

### Anthracycline-SWCNT Binding Kinetics

We next evaluated binding kinetics for individual ssDNA-(*n,m*) combinations using a standard non-cooperative binding model^37^ (**Figure 3, Supplementary Table 2**). Dissociation constants (K_d_) in the sub-micromolar range indicated strong binding affinities of most ssDNA-(*n,m*) constructs for anthracyclines, which is supported by previous studies^40^. We used K_d_ values to identify top-performing ssDNA-*(n,m)* pairings for each anthracycline (**Table 2**). For daunorubicin, intensity modulations of GGACGC-(8,7) exhibited the strongest binding, with a K_d_ of 0.18 µM. Doxorubicin binding to (ATTT)_4_-(10,2) exhibited a K_d_ of 0.13 µM. We observed optimal epirubicin detection with (CTG)_3_C-(9,4), with an intensity change K_d_ of 0.26 µM, while wavelength shifts in CCGCGG-(8,7) were optimal for idarubicin, with a K_d_ of 0.62 µM. In general, average dissociation constants for all anthracyclines were relatively similar, with the average K_d_ for daunorubicin being 7.6 µM, doxorubicin 8.1 µM, and epirubicin and idarubicin being 7.0 and 5.8 µM, respectively. The range of K_d_ values, however, was large. For daunorubicin, K_d_ values ranged from 0.18-60.3 µM, while for doxorubicin and epirubicin, wider K_d_ ranges of 0.13-100.1 µM and 0.26-194.9 µM were seen. Idarubicin responses exhibited a range of K_d_ values between 0.62-74.7 µM. The strong π-π stacking, aromatic binding, and electrostatic interactions between SWCNT and anthracyclines is supported by these relatively strong binding affinities^40, 79-81^. Prior (GT)_15_-SWCNT doxorubicin detection exhibited generally lower binding affinities quantified by intensity changes than those in our study^37^. We also found a wider range of dissociation constants than did prior work with a turn-on doxorubicin-SWCNT sensor^41^.

**Table 2.**
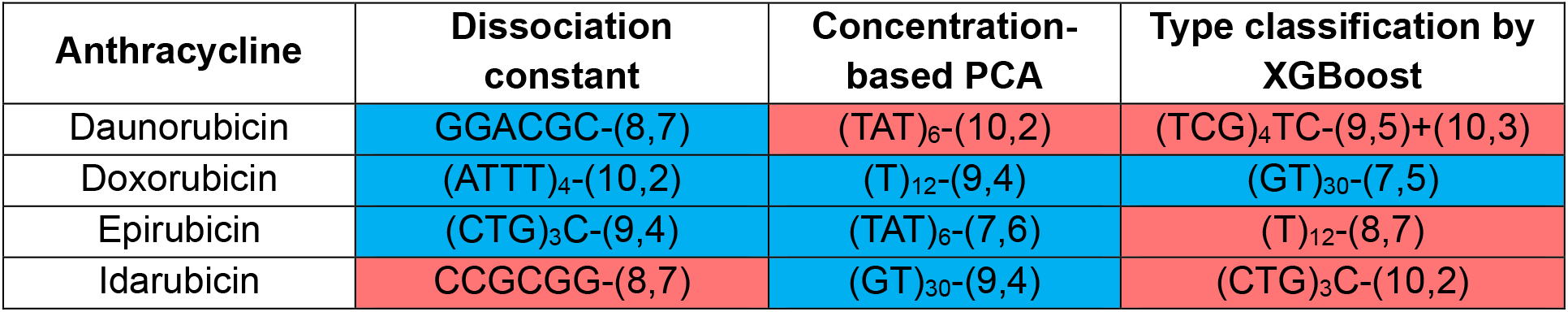
Comparison of ssDNA-*(n,m)* sensor combinations identified to sense anthracyclines. Red cells indicate a wavelength shift as the primary means of detection for the SWCNT, while blue cells indicate a change in intensity.

**Figure 3.**
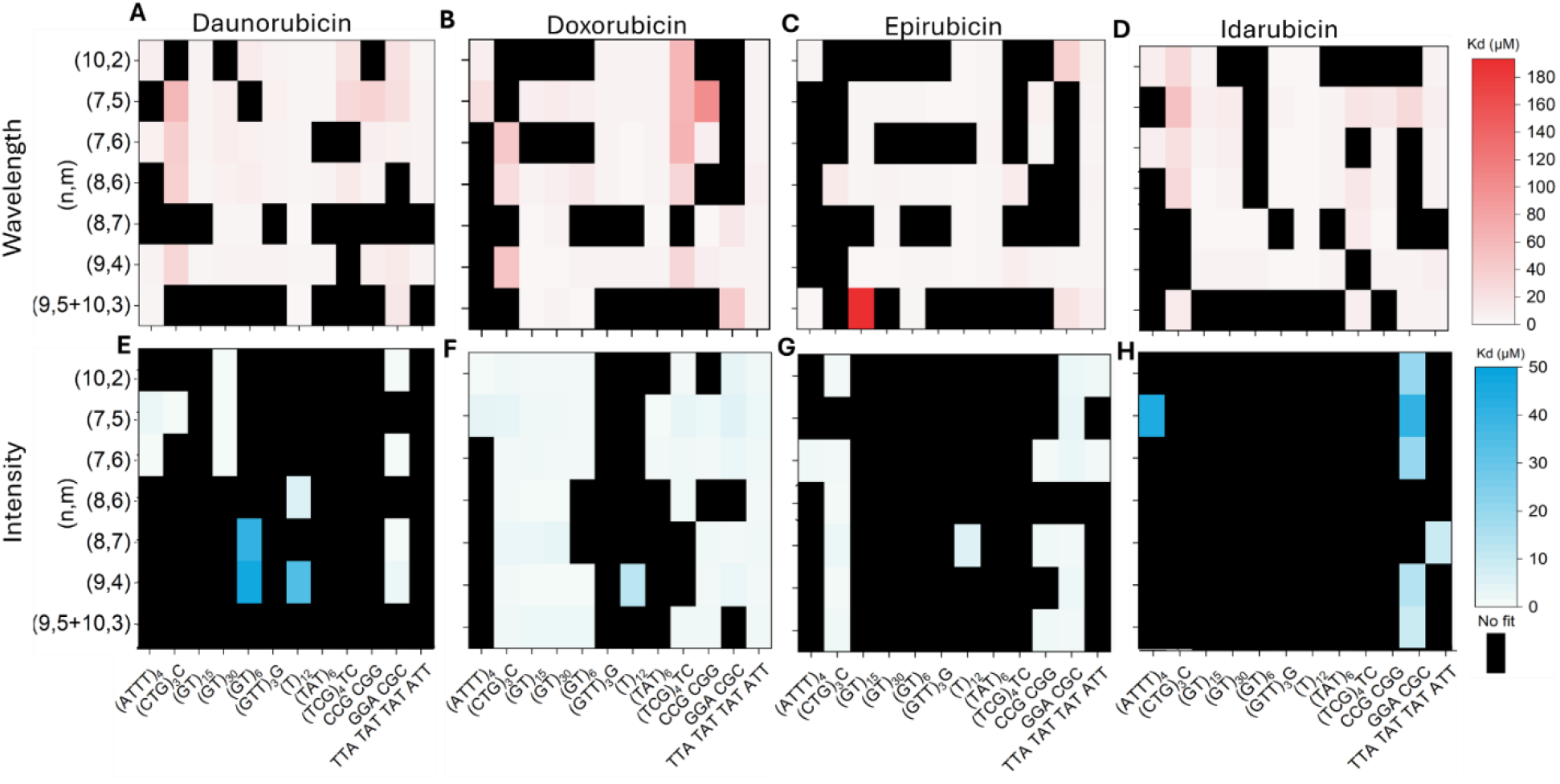
Dissociation constant of each ssDNA-(*n,m*) construct. The Hill equation was used to fit binding kinetics and obtain a dissociation constant (K_d_) for all SWCNT constructs for both wavelength and intensity-induced fluorescence changes in the presence of daunorubicin (**A, E**), doxorubicin (**B**,**F**), epirubicin (**C**,**G**), and idarubicin (**D**,**H**). Black = no fit.

### Statistical Assessment of Significant Sensor Responses

Given different outcomes in optimal ssDNA-(*n,m*) species for LOD and dissociation constant, we next sought to determine which specific features of fluorescence response data best predict the presence of an anthracycline. As the screening experiments yielded 14,112 data points, we used machine learning models to identify spectral fingerprints that improve sensing for each anthracycline (**Figure 1**). First, we ranked dataset features, comprising ssDNA-(*n,m*) combinations and their corresponding spectral responses, using ANOVA. Spectral responses significantly different than controls (*p* < 0.05) at two or more concentration levels for at least one type of anthracycline were included in the feature set. This reduced the feature dimensionality by 57%, yielding 48 statistically significant ssDNA-(*n,m*) combinations with 93 spectral features. The 93 spectral features comprised 45 ssDNA-(*n,m*) constructs exhibiting concurrent changes in both wavelength and intensity, and 3 species characterized by changes in either wavelength or intensity.

The selected nanosensors exhibited diverse response behaviors, with some showing monotonic intensity changes and others displaying wavelength shifts, reflecting heterogeneous ssDNA-(*n,m*) interactions with different target analytes. Interestingly, both wavelength shift and intensity changes demonstrated similar patterns in fluorescence modulations for all ssDNA-*(n,m)* pairings, but the magnitude of change of the spectral signatures was slightly different (**Figure 4**). The most significant features were observed with (CTG)_3_C, which showed 24 significant features for anthracycline and spectral feature pairings. This was followed by TTATATTATATT, (TCG)_4_TC, and (GTT)_3_G which all had 16-18 significant features. Each of these sequences are 10-14 bases, which suggests there may be an optimal length for anthracycline detection. Interestingly, however, (GT)_15_-SWCNT had more significant features among poly-(GT) sequences. (7,5), (7,6), and (9,4) SWCNT were identified as high-contributing features in the ANOVA-based selection, with 32-39 significant features each. This may be due to their relatively high abundance in the HiPCO SWCNT mixture and their on-resonance excitation in these studies. The emission wavelengths of these peaks are between 1040-1137 nm, suggesting that smaller diameter species had more significant changes from perturbations from anthracyclines. However, larger diameter (8,7) and (8,6) exhibited fewer significant features.

**Figure 4.**
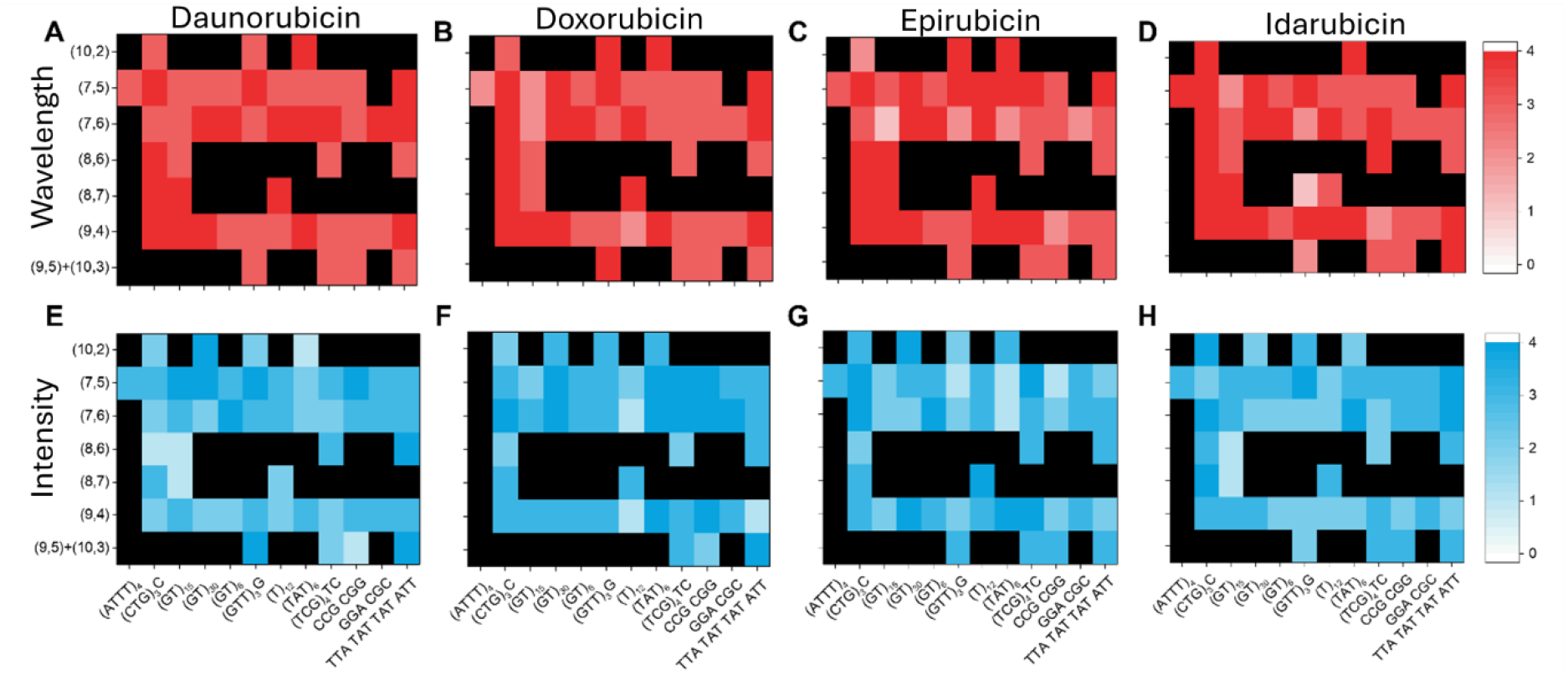
Heatmap plots by ANOVA test for statistically significant feature selection. Wavelength (**A-D**) and intensity (**E-H**) changes of each ssDNA(row) - (*n,m*) (column) combination were correlated with the four concentrations (0.1, 1, 10, and 100 µM) of daunorubicin (**A, E**), doxorubicin (**B**,**F**), epirubicin (**C, G**), and idarubicin (**D, H**). Legend scale indicates the count of significant features among four concentrations.

### Multi-Class Classification of Anthracyclines

The selected feature set of 48 nanosensors was subsequently used as input to PCA. The PCA plot showed the variance in spectral responses across the separate anthracycline classes. While the score plot of the first two principal components (PC1 and PC2) collectively explained 77% of the total variance, the discrimination between anthracycline types was limited, leaving substantial overlap among the four analytes (**Supplementary Figure 8**). This poor separability in PCA indicates the limitations of unsupervised linear dimensionality reduction for anthracycline discrimination, suggesting that more sophisticated machine learning models are needed.

To evaluate classification performance, three machine learning models were implemented: DT, SVM, and XGBoost. The models were trained on spectral responses at 0.1, 1, 10, and 100 µM, and their performance was evaluated on an independent test set with intermediate concentrations of 0.5, 5, and 50 µM. Using 3-fold cross-validation, the models achieved accuracies of 94±12% (DT), 98±4% (SVM), and 100% (XGBoost). Test set performance confirmed these results with accuracies of 100%, 50%, and 100%, respectively (**Figure 5A**).

**Figure 5.**
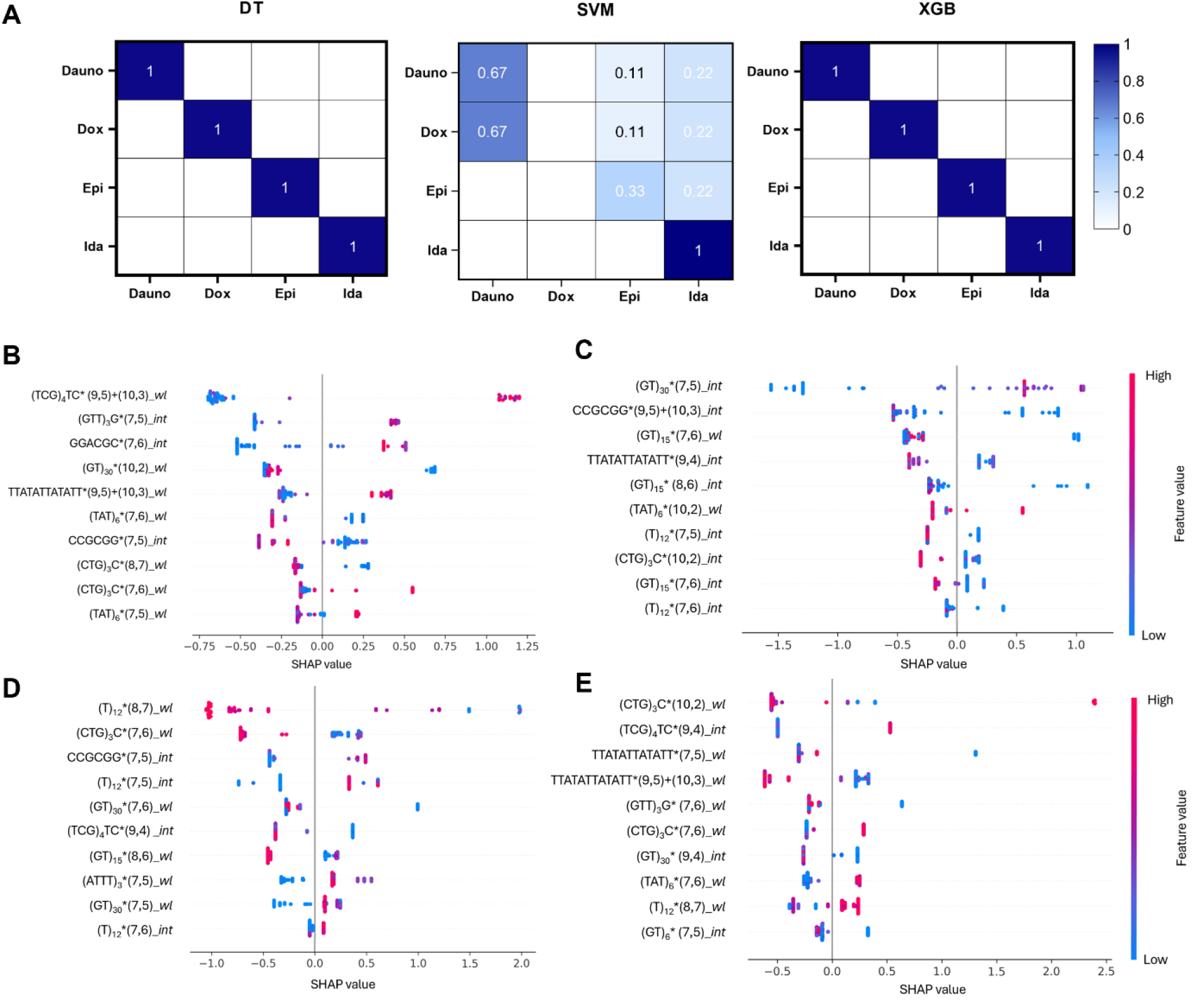
Machine learning to determine optimal features for anthracycline detection. (**A**) Confusion matrix of the three machine learning models for discriminating four anthracycline classes in the independent test set. (**B-E**) SHAP values indicating feature importance for each anthracycline discrimination against others. The colors on the symbols indicate the magnitudes of corresponding input variables, red and blue for higher and lower values, respectively. (**B**) daunorubicin, (**C**) doxorubicin, (**D**) epirubicin, and (**E**) idarubicin.

To define the most discriminable features for anthracycline classification, we calculated SHAP values in the optimized XGB model (**Figure 5 B-E**). By integrating local feature attributions into a unified global interpretability framework, SHAP analysis provided insight into which spectral characteristics most strongly influenced the decision boundaries of the models. Consequently, the derived SHAP values served as an effective means to associate the machine learning predictions with underlying physicochemical variations among the anthracycline molecules. SHAP values for each anthracycline revealed the contribution of each feature to predicting anthracycline type. Features exhibiting the largest absolute SHAP values, such as wavelength shifts for (TCG)_4_TC*-* (9,5)+(10,3) and intensity changes of (GTT)_3_G*-*(7,5), demonstrated the strongest contributing features for distinguishing daunorubicin from other anthracyclines. For doxorubicin, (GT)_30_-(7,5) intensity change and CCGCGG-(9,5)+(10,3) intensity change had the highest SHAP values. The highest-ranking sensor constructs for epirubicin were shifts in (T)_12_-(8,7) and (CTG)_3_C-(7,6). For idarubicin sensing, the top-ranked sensor constructs were wavelength shifts in (CTG)_3_C-(10,2) and intensity change in (TCG)_4_TC-(9,4). These compound-specific classifying features demonstrated that the detection of each anthracycline is characterized by a unique subset of highly informative ssDNA-(*n,m*) combinations that reflect differential binding affinities and sensitivities.

### Concentration-Based Binary Classification

We next investigated whether the sensor array could discriminate variations in anthracycline concentration. First, PCA was performed to visualize concentration-dependent clustering patterns for each anthracycline. By transforming the 93 spectral features into a reduced set of principal components (PCs), major chemical variations were captured within a lower-dimensional space. To further evaluate concentration discrimination capability, PCA was applied to samples for five concentrations (0.1, 1, 5, 10, 100 µM) to establish the principal component space. Subsequently, test samples at intermediate concentrations (0.5 and 5 µM) were projected onto the same principal component space to assess whether they could be reliably assigned to binary concentration groups: low (≤5 µM) or high (>5 µM). The score plots of the first two principal components demonstrate general clustering of low and high concentrations for each anthracycline, although there are some overlaps in doxorubicin and epirubicin **(Figure 6)**. To quantify the accuracy of this clustering in the PCA plot, an SVM was used to find the decision boundaries. Prediction models of daunorubicin and idarubicin achieved 100% accuracy with distinct clustering patterns, whereas doxorubicin and epirubicin demonstrated 50% accuracy with substantial overlap in PC1 and PC2 domains, indicating that their concentration-dependent response patterns are not aligned with linear variance (**Supplementary Table 3**).

**Figure 6.**
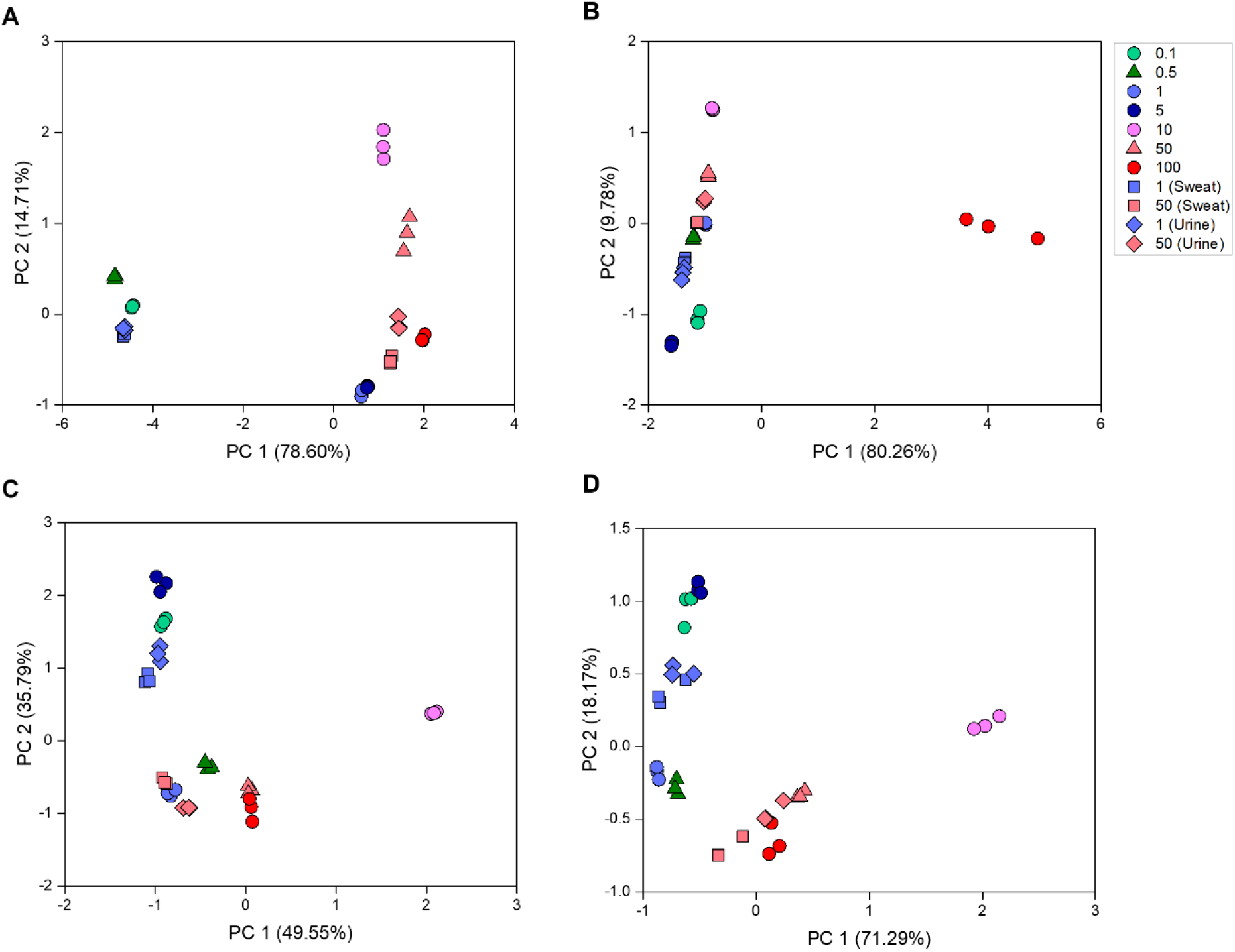
PC score plot for concentration-level discrimination across buffer and biological matrices. PC1 and PC2 domains score plots illustrating the separation of low (≤5 µM) versus high (>5 µM) concentration groups for each anthracycline based on the 49-nanosensor array responses. Training set at defined concentration levels (0.1, 1, 5, 10, 100 µM) is represented by circles, while the test set at intermediate concentration is shown as triangles. Biological matrix samples are indicated by squares (synthetic sweat) and diamonds (synthetic urine). Color denotes concentration: green and blue represent low concentration (≤5 µM) and pink and red represent high concentration (>5 µM) within each anthracycline class. **(A)** daunorubicin, **(B)** doxorubicin, **(C)** epirubicin, and **(D)** idarubicin.

To identify the nanosensor features most responsible for concentration-dependent variance, principal component loadings were analyzed for the first ten principal components (**Supplementary Table 4**). PC loadings quantify the contribution of each feature to the directions of maximum variance, revealing which ssDNA*-(n,m)* combinations are most sensitive to concentration changes within each anthracycline class. For daunorubicin, (TAT)_6_-SWCNT exhibited the highest PC1 loadings in their wavelength response, indicating that wavelength changes from these sensors capture the primary axis of concentration-dependent variance. Doxorubicin classification was dominated by (T)_12_ homopolymer sequences, suggesting that poly-thymine wrapping conformations are particularly sensitive to doxorubicin concentration changes. Epirubicin showed distributed feature importance across multiple sequences, as TTATATTATATT, (GTT)_3_G, and (TAT)_6,_ contributed substantially to PC1 variance, reflecting more heterogeneous sensor response patterns. Idarubicin demonstrated strong sequence dependence on GT-sequences of varying lengths, indicating that GT-rich motifs provide robust recognition. These loadings reveal distinct sequence-dependent recognition patterns for each anthracycline. Strong sensor sequences revealed by machine learning differ from those discovered in a previous array screen for doxorubicin. In that work, AGAATTACTT, flanked by C-rich ends, was used^41^, whereas the machine learning-selected sequences here are either A-T or G-T dominant. Comparing the type of responses observed upon anthracycline interaction, most SWCNTs demonstrated intensity quenching and red shifting across all *(n,m)*, consistent with what has previously been reported for (GT)_15_ based sensors^37, 40^. The preferential sensors identified through machine learning here demonstrated lower dissociation constants for the top selected daunorubicin (K_d_=2.2 µM) and doxorubicin sensors (K_d_= 12.2 µM) compared to the CB-13 sensor (K_d dauno_= 100 µM, K_d dox_ = 14 µM)^41^.

### Sensor Array Performance Validation in Synthetic Sweat and Urine

Synthetic sweat and synthetic urine were used as biofluids in which anthracyclines would be excreted following chemotherapy administration^19, 82, 83^. The same four anthracyclines were spiked into these biological matrices at concentrations of 1 µM and 50 µM for all SWCNTs. To evaluate classification performance, PCA and SVM were applied to predict concentration levels in biological matrices. (**Figure 6**). Daunorubicin and idarubicin achieved perfect binary classification accuracy (100%) in distinguishing low (≤5 µM) versus high (>5 µM) concentration groups across both conditions, demonstrating robust discriminatory capability despite the complex matrix composition. In contrast, doxorubicin and epirubicin exhibited substantially lower accuracy (50%), systematically misclassifying high-concentration samples as low concentration while correctly identifying low-concentration samples (**Supplementary Table 3**). This compound-specific performance discrepancy suggests that doxorubicin and epirubicin generate concentration-dependent response patterns that are either too subtle or non-monotonic to establish reliable decision boundaries using the current feature set and classification algorithm. The differential performance may arise from structural differences in the anthracycline structure—specifically, the C-14 hydroxyl group present in doxorubicin and epirubicin but absent in daunorubicin and idarubicin—which could alter binding modes, electronic coupling to the SWCNT surface, or susceptibility to matrix interference effects^84^. Nevertheless, the robust performance for daunorubicin and idarubicin across complex matrices demonstrates that the nanosensor array maintains clinically relevant discrimination capability for specific anthracyclines in complex biofluids.

### Comparative Analysis to Identify Distinct Optimal Sensors for Each Anthracycline

A comparison of the methods employed for sensor selection revealed no overlap in the preferential sensors (both ssDNA and *(n,m)* species) for each anthracycline (**Table 2**). On the surface, this may be unexpected, however it is good evidence that each model evaluated separate criteria, such as binding affinity, concentration-dependent sensitivity, and other discriminable factors. Interestingly, we observed that larger diameter *(n,m)* species were consistently selected. This finding is supported by prior studies investigating anthracycline adsorption to SWCNT^79, 85^ and molecular dynamics simulations^81^. Above, we noted that smaller diameter SWCNT species contributed more significant features, though we found that this does not translate to optimal detection. While sensor selection based on the dissociation constant has been used before to determine the selectivity and sensitivity of SWCNT sensors^24, 86^, we found that single-construct data were not consistently well-fit, nor were the dissociation constants as low as in other studies. Because of the diverse pool of ssDNA sequences used here, we postulate that a more systematic length or sequence-based screening pool may improve coherence of predictions. However, increased diversity of sequence space may be valuable for improving sensor function and discriminating between similar molecular structures.

## Conclusions

In this work, we designed a SWCNT nanosensor array and demonstrated the potential clinical applications of several optical nanosensors to detect four anthracycline chemotherapeutics. We implemented the use of high-throughput NIR spectroscopy to analyze fluorescence modulations in the presence of a range of concentrations of each anthracycline. We identified specific ssDNA-(*n,m*) constructs that sensitively responded to each anthracycline based on limits of detection and standard non-linear fits of each binding curve. To enhance anthracycline class detection, supervised machine learning models were trained and tested, with XGBoost achieving perfect accuracy (100%) in discriminating all four structurally similar anthracyclines. While anthracyclines may not be co-administered in an individual patient, structurally similar metabolites would be present concurrently as each anthracycline, as well as other small molecule therapeutics administered to a patient. PCA-based and SVM-confirmed concentration discrimination successfully separated daunorubicin and idarubicin but showed poor performance for doxorubicin and epirubicin. Validation in synthetic sweat and urine confirmed robust performance, with daunorubicin and idarubicin maintaining 100% concentration classification accuracy despite matrix effects. It is possible that this sensor development methodology could serve as a framework for small molecule detection for which few or no biorecognition elements exist. Assessment of sensor responses in plasma and whole blood will further demonstrate the clinical relevance of our sensors, allowing physicians to reevaluate patient treatment plans to avoid drug-induced toxicity. Overall, we provide a methodology for small molecule sensor development, which furthers the field of therapeutic drug monitoring by developing a diagnostic tool for the anthracycline drug class and demonstrating the potential of SWCNT-based spectral fingerprinting for such applications.

## Acknowledgements

The authors wish to acknowledge all members of the Williams Lab for discussion and feedback. This work was supported by NIH R35GM142833, The City College of New York Grove School of Engineering, Stony Brook University School of Medicine, and the SUNY Empire Innovation Program #250010 (R.W.). A.R. and A.I. were supported by a G-RISE Ph.D. traineeship from the National Institutes of Health (T32GM136499). M.K. was supported by the NIH (R00-EB033580). This research was funded, in part, by the Advanced Research Projects Agency for Health (ARPA-H) under Agreement No. 1AY2AX000080-01. The views and conclusions contained in this document are those of the authors and should not be interpreted as representing the official policies, either expressed or implied, of the U.S. Government. Y.K. was supported by the International Research & Development Program of the National Research Foundation of Korea (KNRF; RS-2024-00443688).

