## Supplemental Figures for "Optical Spectral Fingerprinting Enables Sensitive Detection of Anthracycline Chemotherapeutics in Synthetic Clinical Biofluids"

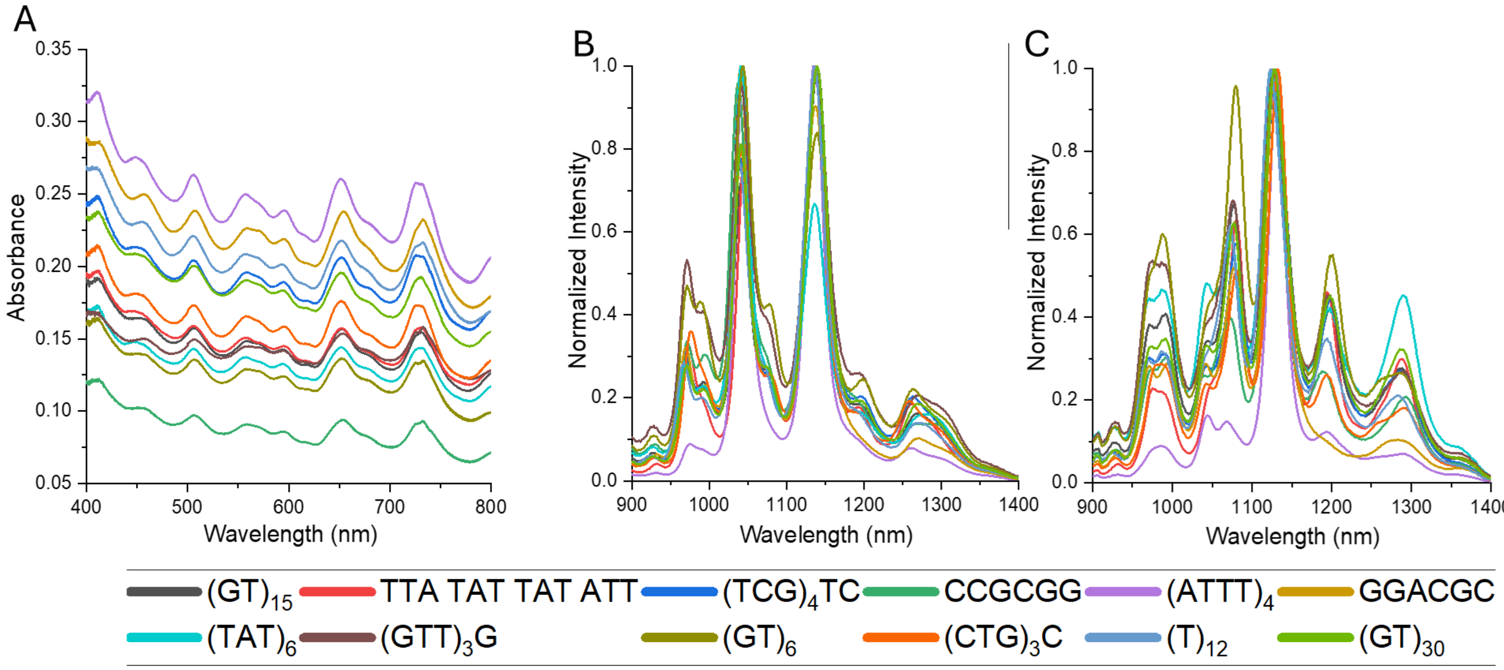


**Supplementary Figure 1**. **Characterization of SWCNT sensor constructs.** A) Representative UV-Vis absorbance spectra. B) Representative fluorescence spectra of SWCNT constructs at excitation wavelength 655 nm. C) Representative fluorescence spectra of SWCNT constructs at excitation wavelength 730 nm.

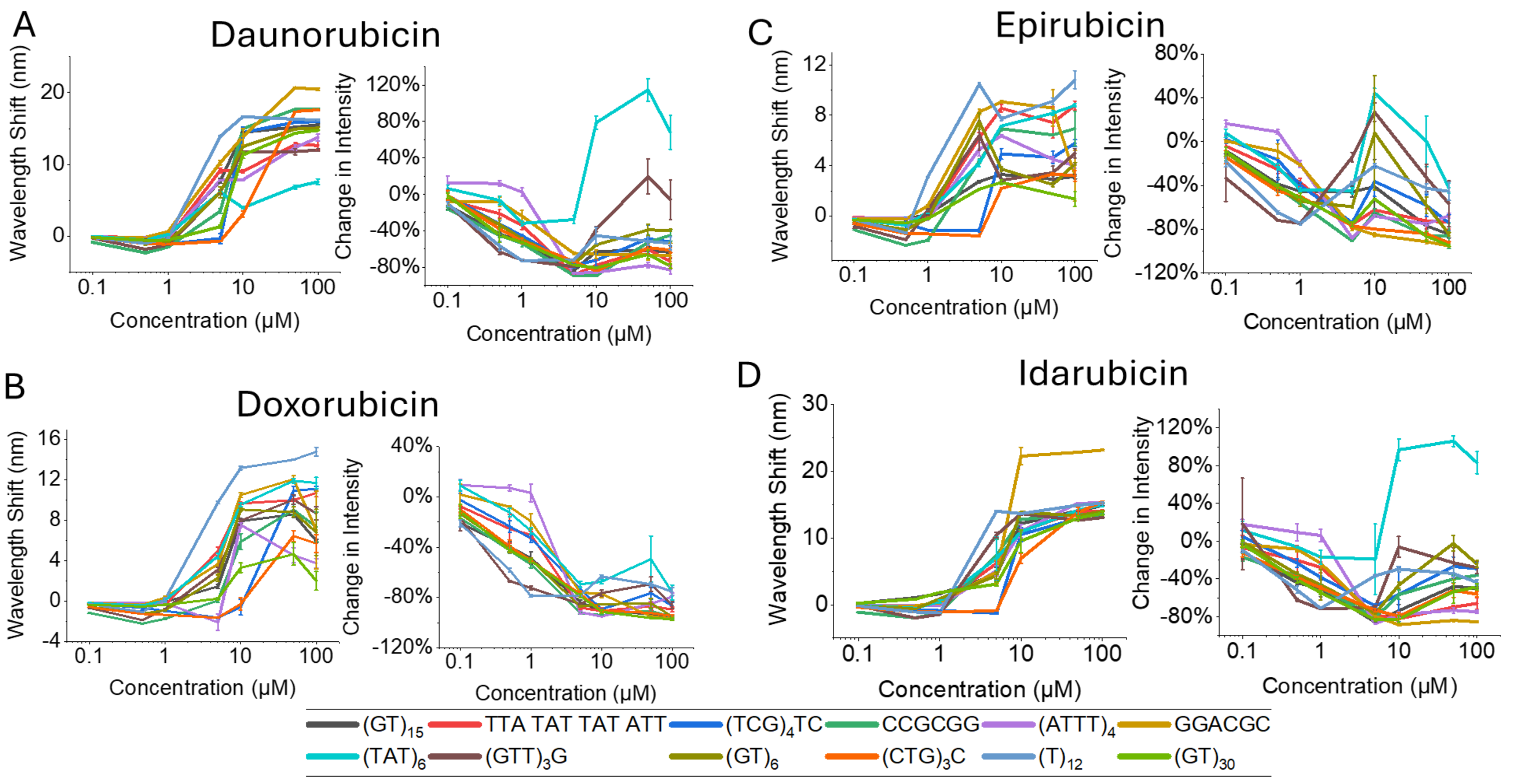


**Supplementary Figure 2.** Concentration response curves of A) daunorubicin, B) doxorubicin, C) epirubicin, and D) idarubicin of all sensor constructs for the (7,6) chirality.

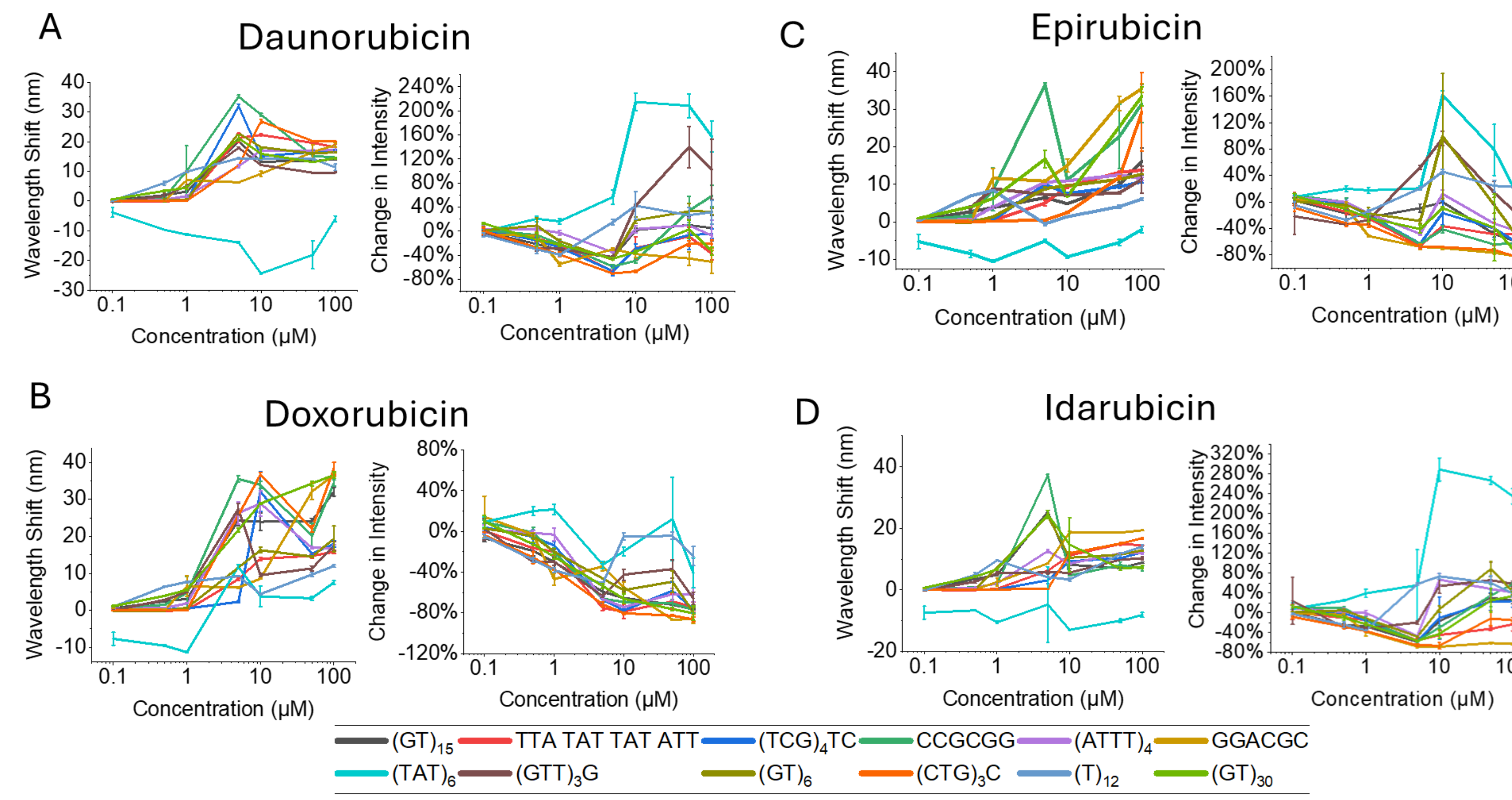


**Supplementary Figure 3.** Concentration response curves of A) daunorubicin, B) doxorubicin, C) epirubicin, and D) idarubicin of all sensor constructs for the (9,5+10,3) chiralities.

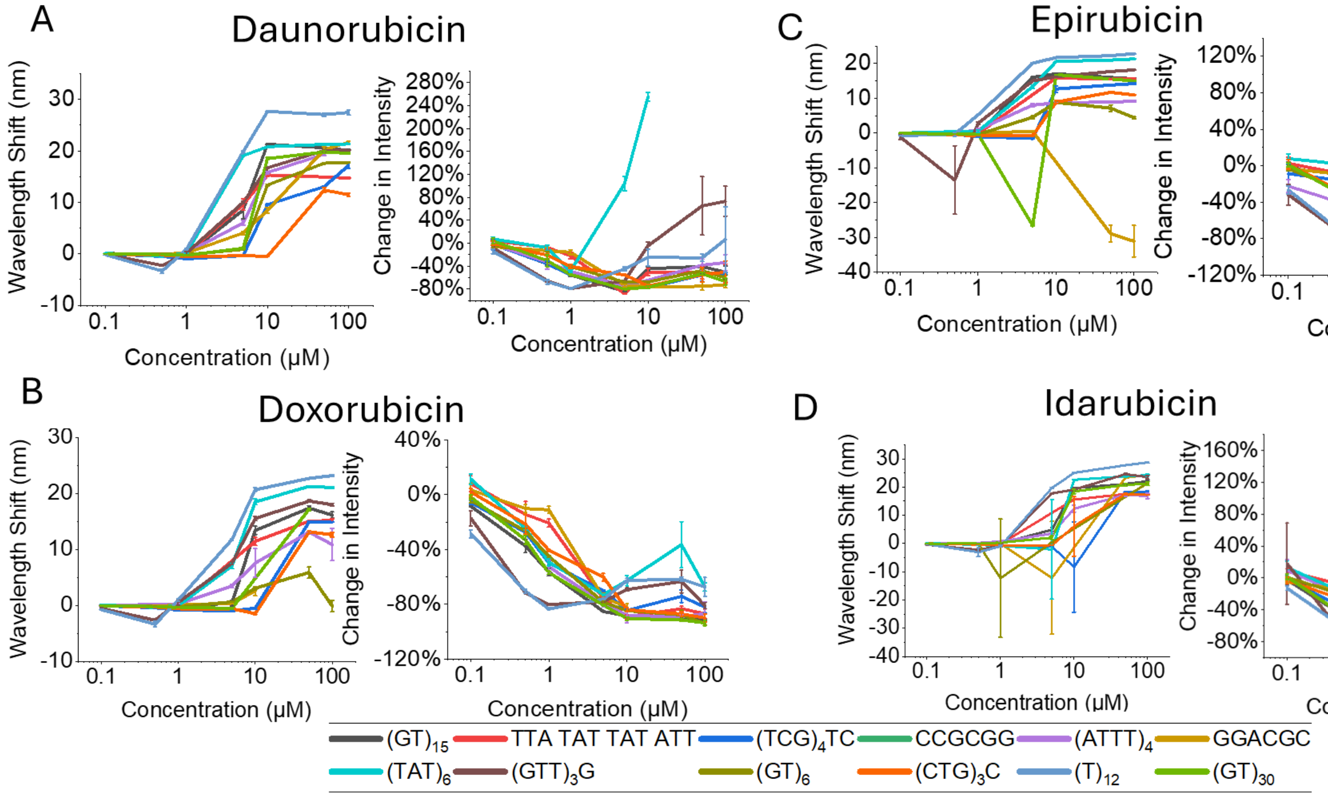


**Supplementary Figure 4.** Concentration response curves of A) daunorubicin, B) doxorubicin, C) epirubicin, and D) idarubicin of all sensor constructs for the (10,2) chirality.

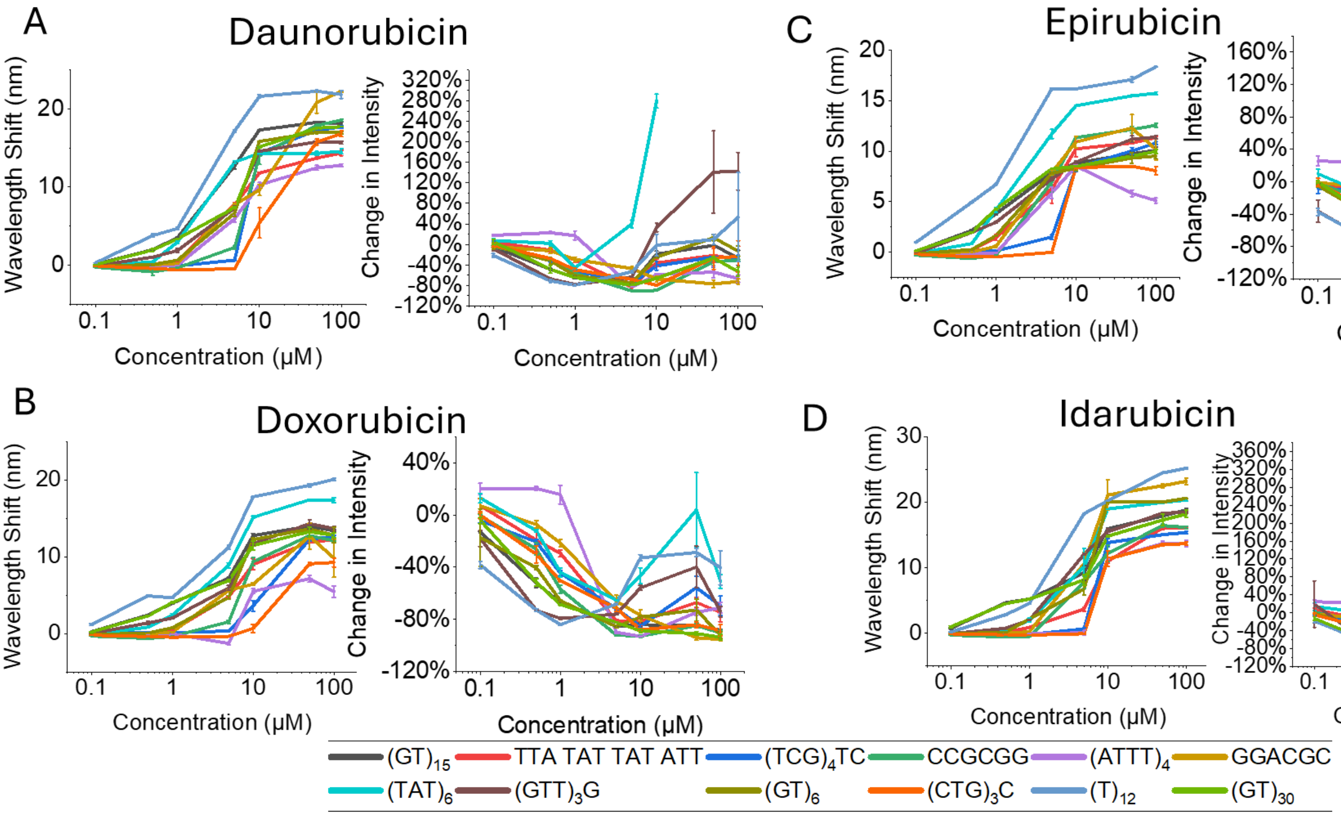


**Supplementary Figure 5.** Concentration response curves of A) daunorubicin, B) doxorubicin, C) epirubicin, and D) idarubicin of all sensor constructs for the (9,4) chirality.

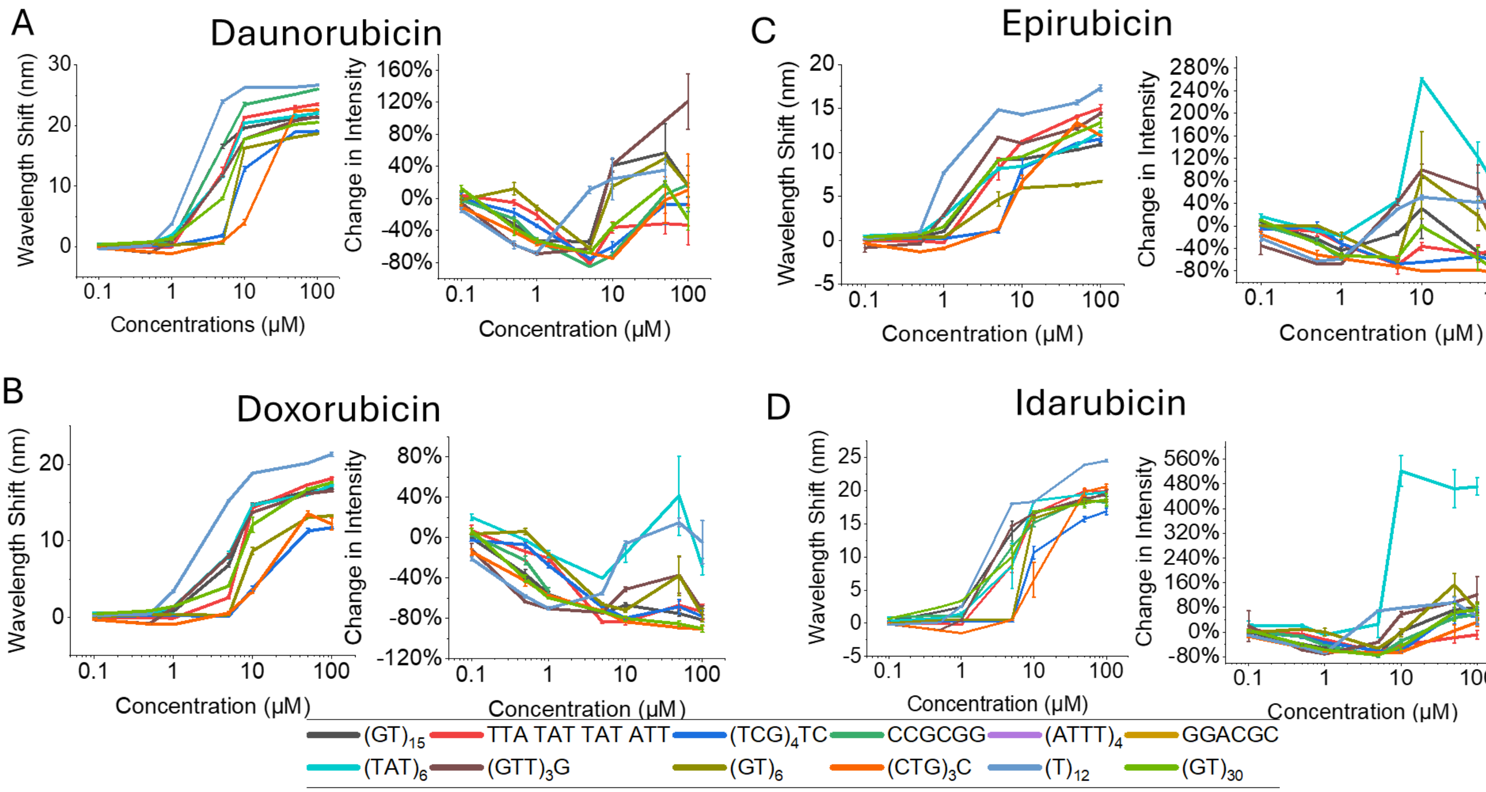


**Supplementary Figure 6.** Concentration response curves of A) daunorubicin, B) doxorubicin, C) epirubicin, and D) idarubicin of all sensor constructs for the (8,6) chirality.

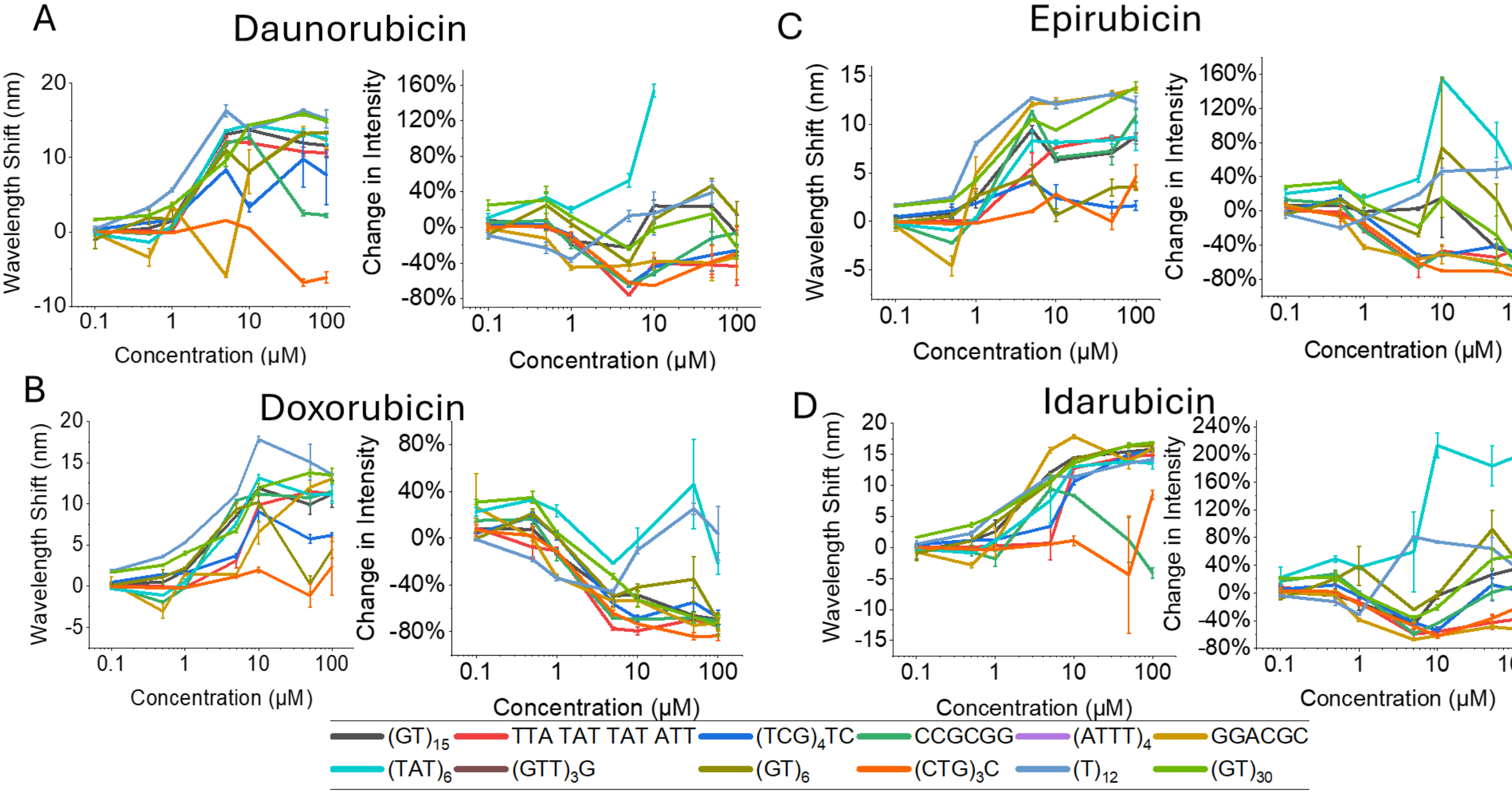


**Supplementary Figure 7.** Concentration response curves of A) daunorubicin, B) doxorubicin, C) epirubicin, and D) idarubicin of all sensor constructs for the (8,7) chirality.

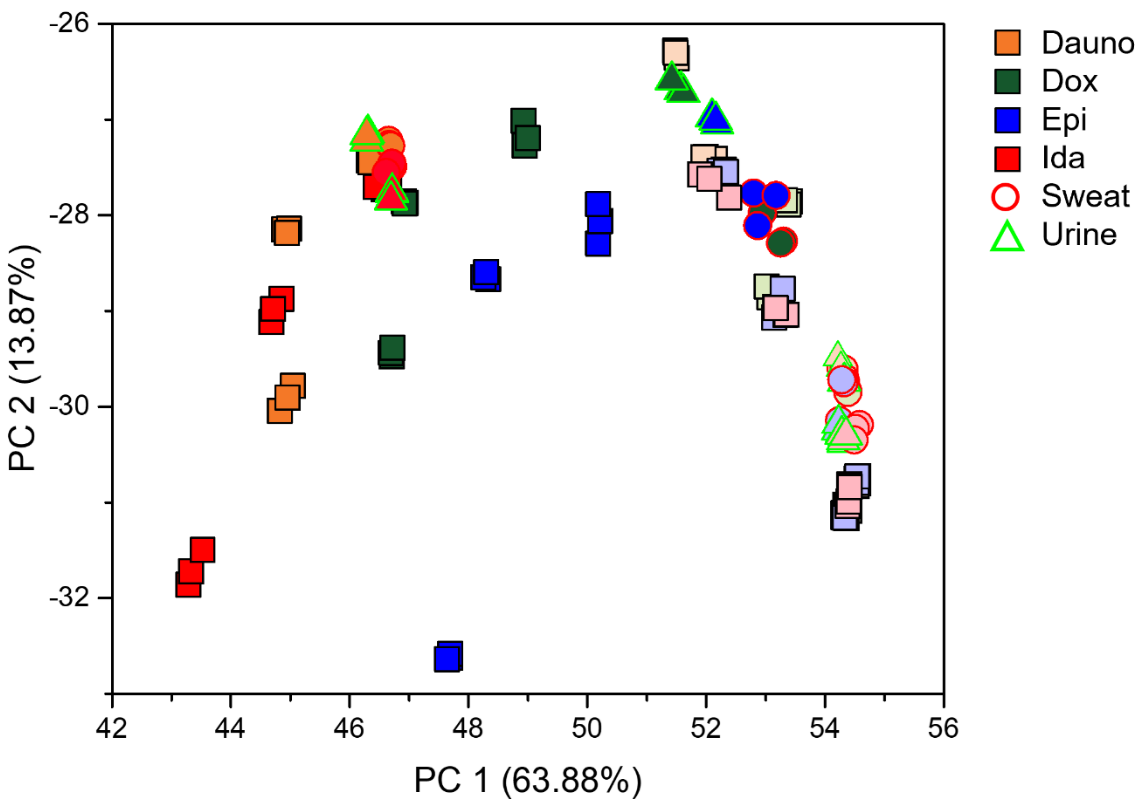


**Supplementary Figure 8** PCA plot of anthracycline classes at different concentrations. Color intensity denotes concentration (bright ≤ 5 μM; dark > 5 μM). Colors indicate anthracycline type: daunorubicin (orange), doxorubicin (green), epirubicin (blue), and idarubicin (red). Shapes denote matrices: buffer (square), synthetic sweat (circle), and synthetic urine (triangle).

**Supplementary Table 1.** Theoretical limits of detection (LOD) (calculated as $3*\sigma_{control}$) for all sensor constructs compared to the experimentally determined LOD (based on ANOVA $p<0.05$).

| Sequence | Chirality | Theoretical LOD_wv | Experimental LOD_wv | Theoretical LOD_int | Experimental LOD_int |
|---|---|---|---|---|---|
| $(GT)_{15}$ | (10,2) | 0.12861 | Dauno - 5<br>Dox – 10<br>Epi - 5<br>Ida - 5 | 0.596 | Dauno – 0.1<br>Dox – 0.1<br>Epi – 0.5<br>Ida - 0.1 |
| $(GT)_{15}$ | (7,5) | 0.3289 | Dauno - 5<br>Dox - 5<br>Epi - 5<br>Ida – 5 | 1.976 | Dauno – 0.1<br>Dox – 0.1<br>Epi – 50<br>Ida – 0.1 |
| $(GT)_{15}$ | (7,6) | 1.39748 | Dauno - 5<br>Dox - 5<br>Epi - 5<br>Ida – 0.5 | 1.634 | Dauno – 0.5<br>Dox – 0.1<br>Epi – 0.5<br>Ida – 0.5 |
| $(GT)_{15}$ | (8,6) | 0.19164 | Dauno – 1 | 0.703 | Dauno – n/a |

| | | | | | |
|---|---|---|---|---|---|
| | | | Dox - 5<br>Epi - 1<br>Ida - 1 | | Dox – 0.1<br>Epi - 50<br>Ida - 100 |
| $(GT)_{15}$ | (8,7) | 3.31338 | Dauno – 5<br>Dox - 5<br>Epi - 5<br>Ida - 5 | 1.151 | Dauno - 100<br>Dox – 0.1<br>Epi - 50<br>Ida – n/a |
| $(GT)_{15}$ | (9,4) | 0.44131 | Dauno – 0.5<br>Dox – 0.5<br>Epi – 0.5<br>Ida – 0.5 | 0.677 | Dauno – 0.5<br>Dox – 0.1<br>Epi - 50<br>Ida – n/a |
| $(GT)_{15}$ | (9,5+10,3) | 0.6772 | Dauno – 0.5<br>Dox - 5<br>Epi - 5<br>Ida – 0.5 | 3.129 | Dauno – 0.1<br>Dox – 0.1<br>Epi – 0.1<br>Ida – 0.1 |
| TTA TAT TAT ATT | (10,2) | 0.06004 | Dauno - 5<br>Dox - 5<br>Epi – n/a<br>Ida - 5 | 1.325 | Dauno - 5<br>Dox - 5<br>Epi – 5<br>Ida – 10 |
| TTA TAT TAT ATT | (7,5) | 0.36057 | Dauno - 5<br>Dox - 5<br>Epi - 5<br>Ida - 5 | 1.183 | Dauno - 5<br>Dox - 5<br>Epi – n/a<br>Ida – n/a |
| TTA TAT TAT ATT | (7,6) | 1.49272 | Dauno - 5<br>Dox - 5<br>Epi - 5<br>Ida - 5 | 1.537 | Dauno - 5<br>Dox - 5<br>Epi - 5<br>Ida - 5 |
| TTA TAT TAT ATT | (8,6) | 0.26255 | Dauno - 5<br>Dox - 5<br>Epi - 5<br>Ida - 5 | 1.54 | Dauno – n/a<br>Dox - 5<br>Epi – n/a<br>Ida – n/a |
| TTA TAT TAT ATT | (8,7) | 0.37601 | Dauno - 5<br>Dox - 5<br>Epi - 5<br>Ida - 10 | 1.514 | Dauno - 100<br>Dox - 5<br>Epi – n/a<br>Ida – n/a |
| TTA TAT TAT ATT | (9,4) | 0.31263 | Dauno - 5<br>Dox - 5<br>Epi - 5<br>Ida - 1 | 1.387 | Dauno – n/a<br>Dox - 5<br>Epi – n/a<br>Ida – n/a |
| TTA TAT TAT ATT | (9,5+10,3) | 0.24238 | Dauno - 5<br>Dox - 5<br>Epi - 5<br>Ida - 5 | 1.849 | Dauno – n/a<br>Dox -5<br>Epi – n/a<br>Ida – n/a |
| $(TCG)_4TC$ | (10,2) | 0.31444 | Dauno – 0.5<br>Dox – 0.5<br>Epi - 1<br>Ida - 1 | 4.235 | Dauno - n/a<br>Dox - n/a<br>Epi - n/a<br>Ida - n/a |

| | | | | | |
|---|---|---|---|---|---|
| $(TCG)_4TC$ | (7,5) | 0.78879 | Dauno - 10<br>Dox - 50<br>Epi - 5<br>Ida - 10 | 4.814 | Dauno – n/a<br>Dox - n/a<br>Epi - n/a<br>Ida - n/a |
| $(TCG)_4TC$ | (7,6) | 0.54083 | Dauno - 10<br>Dox - 50<br>Epi - 10<br>Ida - 10 | 4.878 | Dauno - n/a<br>Dox - n/a<br>Epi - n/a<br>Ida - n/a |
| $(TCG)_4TC$ | (8,6) | 0.10638 | Dauno - 5<br>Dox - 10<br>Epi - 10<br>Ida - 10 | 0.111 | Dauno - n/a<br>Dox - 5<br>Epi - 5<br>Ida - 50 |
| $(TCG)_4TC$ | (8,7) | 0.53573 | Dauno - 50<br>Dox - 1<br>Epi - n/a<br>Ida – 0.5 | 0.103 | Dauno - n/a<br>Dox – 0.1<br>Epi – 0.1<br>Ida - n/a |
| $(TCG)_4TC$ | (9,4) | 0.16406 | Dauno - 5<br>Dox - 10<br>Epi - 5<br>Ida - 10 | 0.108 | Dauno - n/a<br>Dox - 1<br>Epi - n/a<br>Ida - 50 |
| $(TCG)_4TC$ | (9,5+10,3) | 1.63363 | Dauno - 5<br>Dox - 10<br>Epi - 5<br>Ida - 5 | 5.002 | Dauno - n/a<br>Dox - n/a<br>Epi - n/a<br>Ida - n/a |
| CCG CGG | (7,5) | 0.2427 | Dauno - 10<br>Dox - 10<br>Epi – 0.5<br>Ida - 10 | 1.366 | Dauno - n/a<br>Dox - 5<br>Epi - 50<br>Ida - n/a |
| CCG CGG | (7,6) | 0.38861 | Dauno – 0.1<br>Dox - 10<br>Epi – 0.5<br>Ida - 5 | 1.411 | Dauno - n/a<br>Dox - 1<br>Epi - 1<br>Ida - n/a |
| CCG CGG | (8,6) | 0.79 | Dauno - 5<br>Dox - 10<br>Epi – 0.1<br>Ida – 0.1 | 0.2 | Dauno - 1<br>Dox - 10<br>Epi - 1<br>Ida - 50 |
| CCG CGG | (8,7) | 0.56565 | Dauno - 5<br>Dox - 5<br>Epi - 5<br>Ida - 100 | 0.212 | Dauno - 100<br>Dox - 5<br>Epi - 5<br>Ida - 50 |
| CCG CGG | (9,4) | 0.08478 | Dauno - 5<br>Dox - 5<br>Epi - 5<br>Ida - 5 | 0.183 | Dauno – 0.1<br>Dox – 0.1<br>Epi - 1<br>Ida – 0.1 |
| CCG CGG | (9,5+10,3) | 0.9179 | Dauno - 5<br>Dox - 5<br>Epi - 50 | 1.381 | Dauno - 100<br>Dox - 5<br>Epi - n/a |

| | | | | | |
|---|---|---|---|---|---|
| | | | Ida – 0.1 | | Ida - 100 |
| $(ATTT)_4$ | (10,2) | 0.31884 | Dauno – 0.1<br>Dox - 5<br>Epi – 0.1<br>Ida – 0.1 | 1.506 | Dauno - n/a<br>Dox - n/a<br>Epi - n/a<br>Ida - n/a |
| $(ATTT)_4$ | (7,5) | 0.26073 | Dauno - 10<br>Dox - 10<br>Epi - 5<br>Ida - 10 | 0.896 | Dauno - 5<br>Dox - 10<br>Epi - 50<br>Ida - 10 |
| $(ATTT)_4$ | (9,5+10,3) | 0.21447 | Dauno - 1<br>Dox - 5<br>Epi – 0.5<br>Ida – 0.5 | 1.064 | Dauno - n/a<br>Dox - n/a<br>Epi - n/a<br>Ida - 10 |
| GGA CGC | (10,2) | 0.17449 | Dauno - 5<br>Dox - 10<br>Epi - 50<br>Ida - 5 | 1.472 | Dauno - 5<br>Dox - 5<br>Epi - 50<br>Ida - 5 |
| GGA CGC | (7,5) | 0.41245 | Dauno - 5<br>Dox - 10<br>Epi - 5<br>Ida - 10 | 2.066 | Dauno - n/a<br>Dox - 50<br>Epi - 50<br>Ida - n/a |
| GGA CGC | (7,6) | 1.21187 | Dauno – 5<br>Dox – 10<br>Epi - 5<br>Ida - 5 | 2.128 | Dauno - n/a<br>Dox - 50<br>Epi - 50<br>Ida - n/a |
| GGA CGC | (8,7) | 0.27673 | Dauno - 10<br>Dox - 1<br>Epi - 1<br>Ida - 1 | 0.613 | Dauno - n/a<br>Dox - 50<br>Epi - 50<br>Ida - 100 |
| GGA CGC | (9,4) | 0.21677 | Dauno - 5<br>Dox - 5<br>Epi - 5<br>Ida - 5 | 1.51 | Dauno - 10<br>Dox - 5<br>Epi - 5<br>Ida - 5 |
| GGA CGC | (9,5+10,3) | 1.24989 | Dauno - 1<br>Dox - 1<br>Epi - 1<br>Ida - 1 | 2.587 | Dauno - n/a<br>Dox - n/a<br>Epi - n/a<br>Ida - n/a |
| $(TAT)_6$ | (10,2) | 0.08146 | Dauno - 5<br>Dox - 5<br>Epi - 5<br>Ida – 0.5 | 0.5 | Dauno - 10<br>Dox - 1<br>Epi - n/a<br>Ida - 10 |
| $(TAT)_6$ | (7,5) | 0.01726 | Dauno - 1<br>Dox - 1<br>Epi – 0.5<br>Ida - 5 | 0.716 | Dauno - 10<br>Dox – 0.5<br>Epi - 10<br>Ida - 10 |
| $(TAT)_6$ | (7,6) | 0.14425 | Dauno - 5<br>Dox - 5 | 0.728 | Dauno - 10<br>Dox - 5 |

| | | | | | |
|---|---|---|---|---|---|
| | | | Epi – 0.5<br>Ida - 5 | | Epi - 1<br>Ida - 10 |
| $(TAT)_6$ | (8,6) | 0.31734 | Dauno - 1<br>Dox - 1<br>Epi - 1<br>Ida - 5 | 0.536 | Dauno - 10<br>Dox - 100<br>Epi - 5<br>Ida - 10 |
| $(TAT)_6$ | (8,7) | 1.69853 | Dauno - 1<br>Dox - 1<br>Epi - 5<br>Ida - 5 | 0.57 | Dauno - 10<br>Dox - n/a<br>Epi - 5<br>Ida - 10 |
| $(TAT)_6$ | (9,4) | 0.27307 | Dauno - 1<br>Dox – 0.5<br>Epi – 0.5<br>Ida - 5 | 0.565 | Dauno - 10<br>Dox - n/a<br>Epi - n/a<br>Ida - 10 |
| $(TAT)_6$ | (9,5+10,3) | 7.44552 | Dauno - n/a<br>Dox - 1<br>Epi - n/a<br>Ida - n/a | 0.23 | Dauno - 5<br>Dox - n/a<br>Epi - n/a<br>Ida - 10 |
| $(GTT)_3G$ | (10,2) | 1.48603 | Dauno - 5<br>Dox - 5<br>Epi - 5<br>Ida - 5 | 0.49 | Dauno - n/a<br>Dox – 0.1<br>Epi - 100<br>Ida - n/a |
| $(GTT)_3G$ | (7,5) | 0.96204 | Dauno - 1<br>Dox - 1<br>Epi - 1<br>Ida - 1 | 0.6 | Dauno - 50<br>Dox - 100<br>Epi - n/a<br>Ida - 10 |
| $(GTT)_3G$ | (7,6) | 0.46275 | Dauno - 5<br>Dox - 5<br>Epi - 1<br>Ida - 5 | 0.503 | Dauno - 50<br>Dox - 100<br>Epi - n/a<br>Ida - n/a |
| $(GTT)_3G$ | (8,6) | 0.06248 | Dauno – 0.5<br>Dox – 0.5<br>Epi - 1<br>Ida - 5 | 0.403 | Dauno - 50<br>Dox – 0.5<br>Epi - n/a<br>Ida - 50 |
| $(GTT)_3G$ | (9,4) | 0.02869 | Dauno – 0.5<br>Dox – 0.5<br>Epi – 0.1<br>Ida - 1 | 0.479 | Dauno - 50<br>Dox – 0.5<br>Epi - n/a<br>Ida - n/a |
| $(GTT)_3G$ | (9,5+10,3) | 2.79266 | Dauno – 0.5<br>Dox – 0.5<br>Epi – 0.5<br>Ida – 0.5 | 0.466 | Dauno - 10<br>Dox - n/a<br>Epi - n/a<br>Ida - 10 |
| $(GT)_6$ | (10,2) | 0.44529 | Dauno - 1<br>Dox - n/a<br>Epi - 5<br>Ida - 100 | 0.999 | Dauno - 100<br>Dox - 5<br>Epi - 100<br>Ida - n/a |
| $(GT)_6$ | (7,5) | 0.97191 | Dauno - 5 | 0.977 | Dauno - n/a |

| | | | | | |
|---|---|---|---|---|---|
| | | | Dox - 5<br>Epi - 1<br>Ida - 5 | | Dox - 5<br>Epi - n/a<br>Ida - n/a |
| $(GT)_6$ | (7,6) | 0.40084 | Dauno - 5<br>Dox - 5<br>Epi - 5<br>Ida - 1 | 0.954 | Dauno - n/a<br>Dox - 5<br>Epi - n/a<br>Ida - n/a |
| $(GT)_6$ | (8,6) | 0.39931 | Dauno - 5<br>Dox - 10<br>Epi - 5<br>Ida - 10 | 1.065 | Dauno - 50<br>Dox - 100<br>Epi - n/a<br>Ida - 50 |
| $(GT)_6$ | (8,7) | 1.86924 | Dauno - 5<br>Dox - 100<br>Epi - 50<br>Ida - 1 | 0.267 | Dauno - 10<br>Dox - 100<br>Epi - n/a<br>Ida - 50 |
| $(GT)_6$ | (9,4) | 0.62225 | Dauno - 1<br>Dox - 5<br>Epi - 1<br>Ida - 1 | 1.012 | Dauno - n/a<br>Dox - 1<br>Epi - n/a<br>Ida - n/a |
| $(GT)_6$ | (9,5+10,3) | 1.22519 | Dauno - 5<br>Dox - 10<br>Epi - 1<br>Ida - 5 | 0.886 | Dauno - 10<br>Dox - 100<br>Epi - n/a<br>Ida - 10 |
| $(CTG)_3C$ | (10,2) | 0.08326 | Dauno - 50<br>Dox - 1<br>Epi – 0.5<br>Ida - 50 | 0.25 | Dauno - 5<br>Dox - 1<br>Epi - 1<br>Ida - 1 |
| $(CTG)_3C$ | (7,5) | 0.0253 | Dauno - 50<br>Dox - 50<br>Epi - 10<br>Ida - 50 | 0.11 | Dauno - 1<br>Dox - 5<br>Epi - 5<br>Ida - 100 |
| $(CTG)_3C$ | (7,6) | 0.10945 | Dauno – 0.5<br>Dox - 50<br>Epi – 0.1<br>Ida - 10 | 0.104 | Dauno – 0.1<br>Dox – 0.1<br>Epi – 0.5<br>Ida – 0.5 |
| $(CTG)_3C$ | (8,6) | 0.3439 | Dauno – 0.5<br>Dox - 10<br>Epi – 0.5<br>Ida - 10 | 0.189 | Dauno - n/a<br>Dox – 0.5<br>Epi – 0.5<br>Ida - 100 |
| $(CTG)_3C$ | (8,7) | 1.41223 | Dauno - 50<br>Dox - n/a<br>Epi - 100<br>Ida - n/a | 0.173 | Dauno - n/a<br>Dox - 5<br>Epi - 5<br>Ida - n/a |
| $(CTG)_3C$ | (9,4) | 0.13168 | Dauno - 10<br>Dox - 50<br>Epi - 10<br>Ida - 10 | 0.241 | Dauno - n/a<br>Dox – 0.1<br>Epi – 0.5<br>Ida - n/a |

| | | | | | |
|---|---|---|---|---|---|
| $(CTG)_3C$ | (9,5+10,3) | 0.10956 | Dauno - 5<br>Dox - 10<br>Epi - 100<br>Ida - 10 | 0.117 | Dauno - n/a<br>Dox – 0.1<br>Epi – 0.5<br>Ida - n/a |
| $(T)_{12}$ | (10,2) | 0.4144 | Dauno - 5<br>Dox – 0.1<br>Epi - 1<br>Ida - 5 | 1.447 | Dauno - n/a<br>Dox – 0.5<br>Epi - 100<br>Ida - n/a |
| $(T)_{12}$ | (7,5) | 0.18774 | Dauno – 0.5<br>Dox - 1<br>Epi – 0.5<br>Ida - 1 | 0.239 | Dauno - n/a<br>Dox – 0.5<br>Epi - n/a<br>Ida - n/a |
| $(T)_{12}$ | (7,6) | 0.53042 | Dauno - 5<br>Dox - 5<br>Epi – 0.5<br>Ida – 0.5 | 2.643 | Dauno - n/a<br>Dox - n/a<br>Epi - n/a<br>Ida - n/a |
| $(T)_{12}$ | (8,6) | 0.50043 | Dauno - 1<br>Dox - 1<br>Epi – 0.5<br>Ida - 1 | 1.501 | Dauno - n/a<br>Dox - n/a<br>Epi - n/a<br>Ida - n/a |
| $(T)_{12}$ | (8,7) | 4.20116 | Dauno – 0.5<br>Dox – 0.5<br>Epi – 0.5<br>Ida – 0.5 | 1.527 | Dauno - n/a<br>Dox - n/a<br>Epi - n/a<br>Ida - n/a |
| $(T)_{12}$ | (9,4) | 0.16061 | Dauno – 0.5<br>Dox – 0.1<br>Epi – 0.1<br>Ida – 0.5 | 1.509 | Dauno - n/a<br>Dox - 100<br>Epi - n/a<br>Ida - n/a |
| $(T)_{12}$ | (9,5+10,3) | 1.53304 | Dauno – 0.5<br>Dox – 0.1<br>Epi - 10<br>Ida – 0.5 | 2.771 | Dauno - n/a<br>Dox - n/a<br>Epi - n/a<br>Ida - n/a |
| $(GT)_{30}$ | (10,2) | 0.16727 | Dauno - 5<br>Dox - n/a<br>Epi – 0.5<br>Ida - 5 | ##### | Dauno - 1<br>Dox - 1<br>Epi - 50<br>Ida - n/a |
| $(GT)_{30}$ | (7,5) | 0.08858 | Dauno - 5<br>Dox - 5<br>Epi – 0.5<br>Ida - 10 | 0.452 | Dauno - 5<br>Dox - 1<br>Epi - 10<br>Ida - n/a |
| $(GT)_{30}$ | (7,6) | 0.78338 | Dauno - 5<br>Dox - 10<br>Epi - 5<br>Ida - 5 | 0.5 | Dauno – 0.1<br>Dox – 0.5<br>Epi - 1<br>Ida - 1 |
| $(GT)_{30}$ | (8,6) | 0.27241 | Dauno – 0.5<br>Dox – 5<br>Epi – 0.5 | 0.31 | Dauno - n/a<br>Dox – 0.1<br>Epi - 1 |

| | | | | | |
|---|---|---|---|---|---|
| | | | Ida – 0.5 | | Ida - 50 |
| $(GT)_{30}$ | (8,7) | 3.62661 | Dauno – 0.1<br>Dox – 0.1<br>Epi – 0.1<br>Ida – 0.1 | 0.313 | Dauno - n/a<br>Dox - 10<br>Epi - 10<br>Ida - 50 |
| $(GT)_{30}$ | (9,4) | 0.07942 | Dauno – 0.5<br>Dox – 0.5<br>Epi – 0.5<br>Ida – 0.5 | 0.31 | Dauno - 100<br>Dox – 0.1<br>Epi - 50<br>Ida - n/a |
| $(GT)_{30}$ | (9,5+10,3) | 0.83466 | Dauno – 0.5<br>Dox – 0.5<br>Epi - 1<br>Ida - n/a | 0.54 | Dauno - n/a<br>Dox - 5<br>Epi - 50<br>Ida - 50 |

**Supplementary Table 2.** Logistic model results of wavelength and intensity-induced anthracycline responses

| Sequence | Anthracycline | ***(n,m)*** species | $K_d$_wv | $K_d$_int |
|---|---|---|---|---|
| $(GT)_{15}$ | Dauno | (10,2) | 4.084 | Fit failed |
| $(GT)_{15}$ | Dauno | (7,5) | 4.223 | Fit failed |
| $(GT)_{15}$ | Dauno | (7,6) | 3.965 | 0.182 |
| $(GT)_{15}$ | Dauno | (8,6) | 2.566 | Fit failed |
| $(GT)_{15}$ | Dauno | (8,7) | 1.455 | Fit failed |
| $(GT)_{15}$ | Dauno | (9,4) | 2.375 | Fit failed |
| $(GT)_{15}$ | Dauno | (9,5+10,3) | Fit failed | Fit failed |
| $(GT)_{15}$ | Dox | (10,2) | 10.21 | 0.583 |
| $(GT)_{15}$ | Dox | (7,5) | 7.394 | 1.477 |
| $(GT)_{15}$ | Dox | (7,6) | 4.818 | 1.206 |
| $(GT)_{15}$ | Dox | (8,6) | 5.058 | 0.307 |
| $(GT)_{15}$ | Dox | (8,7) | 1.981 | 2.268 |
| $(GT)_{15}$ | Dox | (9,4) | 3.166 | 0.247 |
| $(GT)_{15}$ | Dox | (9,5+10,3) | 2.314 | 1.616 |
| $(GT)_{15}$ | Epi | (10,2) | 1.906 | Fit failed |
| $(GT)_{15}$ | Epi | (7,5) | 2.027 | Fit failed |
| $(GT)_{15}$ | Epi | (7,6) | 1.937 | Fit failed |
| $(GT)_{15}$ | Epi | (8,6) | 2.25 | Fit failed |
| $(GT)_{15}$ | Epi | (8,7) | 1.256 | 108 |
| $(GT)_{15}$ | Epi | (9,4) | 1.416 | Fit failed |
| $(GT)_{15}$ | Epi | (9,5+10,3) | 194.9 | Fit failed |
| $(GT)_{15}$ | Ida | (10,2) | 6.619 | Fit failed |
| $(GT)_{15}$ | Ida | (7,5) | 6.871 | Fit failed |
| $(GT)_{15}$ | Ida | (7,6) | 6.631 | Fit failed |
| $(GT)_{15}$ | Ida | (8,6) | 2.721 | Fit failed |

| | | | | |
|---|---|---|---|---|
| $(GT)_{15}$ | Ida | (8,7) | 2.071 | Fit failed |
| $(GT)_{15}$ | Ida | (9,4) | 4.249 | Fit failed |
| $(GT)_{15}$ | Ida | (9,5+10,3) | Fit failed | Fit failed |
| TTA TAT TAT ATT | Dauno | (10,2) | 2.932 | Fit failed |
| TTA TAT TAT ATT | Dauno | (7,5) | 3.349 | Fit failed |
| TTA TAT TAT ATT | Dauno | (7,6) | 3.524 | 0.75 |
| TTA TAT TAT ATT | Dauno | (8,6) | 3.911 | Fit failed |
| TTA TAT TAT ATT | Dauno | (8,7) | 1.664 | Fit failed |
| TTA TAT TAT ATT | Dauno | (9,4) | 4.047 | Fit failed |
| TTA TAT TAT ATT | Dauno | (9,5+10,3) | 1.643 | Fit failed |
| TTA TAT TAT ATT | Dox | (10,2) | 5.11 | 1.303 |
| TTA TAT TAT ATT | Dox | (7,5) | 4.914 | 1.53 |
| TTA TAT TAT ATT | Dox | (7,6) | 4.223 | 1.255 |
| TTA TAT TAT ATT | Dox | (8,6) | 8.85 | 0.93 |
| TTA TAT TAT ATT | Dox | (8,7) | 6.372 | 1.266 |
| TTA TAT TAT ATT | Dox | (9,4) | 6.245 | 0.727 |
| TTA TAT TAT ATT | Dox | (9,5+10,3) | 3.795 | 1.174 |
| TTA TAT TAT ATT | Epi | (10,2) | 5.11 | 0.863 |
| TTA TAT TAT ATT | Epi | (7,5) | 3.945 | Fit failed |
| TTA TAT TAT ATT | Epi | (7,6) | 2.512 | 0.738 |
| TTA TAT TAT ATT | Epi | (8,6) | 4.483 | Fit failed |
| TTA TAT TAT ATT | Epi | (8,7) | 3.307 | Fit failed |
| TTA TAT TAT ATT | Epi | (9,4) | 3.33 | 0.277 |
| TTA TAT TAT ATT | Epi | (9,5+10,3) | 7.184 | 0.507 |
| TTA TAT TAT ATT | Ida | (10,2) | 3.748 | 1.278 |
| TTA TAT TAT ATT | Ida | (7,5) | 9.713 | 1.02 |
| TTA TAT TAT ATT | Ida | (7,6) | 5.254 | 0.981 |
| TTA TAT TAT ATT | Ida | (8,6) | 5.074 | Fit failed |
| TTA TAT TAT ATT | Ida | (8,7) | 9.396 | 0.796 |
| TTA TAT TAT ATT | Ida | (9,4) | 9.605 | Fit failed |

| | | | | |
|---|---|---|---|---|
| TTA TAT TAT ATT | Ida | (9,5+10,3) | 5.672 | Fit failed |
| $(TCG)_4TC$ | Dauno | (10,2) | 18.47 | Fit failed |
| $(TCG)_4TC$ | Dauno | (7,5) | 29.48 | 0.834 |
| $(TCG)_4TC$ | Dauno | (7,6) | 9.18 | Fit failed |
| $(TCG)_4TC$ | Dauno | (8,6) | 12.21 | Fit failed |
| $(TCG)_4TC$ | Dauno | (8,7) | Fit failed | Fit failed |
| $(TCG)_4TC$ | Dauno | (9,4) | 10.2 | Fit failed |
| $(TCG)_4TC$ | Dauno | (9,5+10,3) | Fit failed | Fit failed |
| $(TCG)_4TC$ | Dox | (10,2) | 60.9 | 0.649 |
| $(TCG)_4TC$ | Dox | (7,5) | 60.61 | 2.82 |
| $(TCG)_4TC$ | Dox | (7,6) | 64.27 | 1.1 |
| $(TCG)_4TC$ | Dox | (8,6) | 32.17 | 1.322 |
| $(TCG)_4TC$ | Dox | (8,7) | Fit failed | 2.079 |
| $(TCG)_4TC$ | Dox | (9,4) | 32.89 | 0.562 |
| $(TCG)_4TC$ | Dox | (9,5+10,3) | Fit failed | 1.394 |
| $(TCG)_4TC$ | Epi | (10,2) | 10.66 | Fit failed |
| $(TCG)_4TC$ | Epi | (7,5) | 8.839 | Fit failed |
| $(TCG)_4TC$ | Epi | (7,6) | Fit failed | Fit failed |
| $(TCG)_4TC$ | Epi | (8,6) | 11.43 | 1.021 |
| $(TCG)_4TC$ | Epi | (8,7) | Fit failed | Fit failed |
| $(TCG)_4TC$ | Epi | (9,4) | 9.134 | Fit failed |
| $(TCG)_4TC$ | Epi | (9,5+10,3) | 2.141 | Fit failed |
| $(TCG)_4TC$ | Ida | (10,2) | Fit failed | Fit failed |
| $(TCG)_4TC$ | Ida | (7,5) | 19.24 | Fit failed |
| $(TCG)_4TC$ | Ida | (7,6) | 12.65 | Fit failed |
| $(TCG)_4TC$ | Ida | (8,6) | 15.2 | Fit failed |

| | | | | |
|---|---|---|---|---|
| $(TCG)_4TC$ | Ida | (8,7) | 10.64 | Fit failed |
| $(TCG)_4TC$ | Ida | (9,4) | 8.752 | Fit failed |
| $((TCG)_4TC$ | Ida | (9,5+10,3) | 7.325 | Fit failed |
| CCG CGG | Dauno | (10,2) | Fit failed | Fit failed |
| CCG CGG | Dauno | (7,5) | 34.76 | Fit failed |
| CCG CGG | Dauno | (7,6) | 6.803 | Fit failed |
| CCG CGG | Dauno | (8,6) | 3.536 | Fit failed |
| CCG CGG | Dauno | (8,7) | Fit failed | Fit failed |
| CCG CGG | Dauno | (9,4) | 9.385 | Fit failed |
| CCG CGG | Dauno | (9,5+10,3) | Fit failed | Fit failed |
| CCG CGG | Dox | (10,2) | Fit failed | Fit failed |
| CCG CGG | Dox | (7,5) | 100.1 | 1.599 |
| CCG CGG | Dox | (7,6) | 8.531 | 0.75 |
| CCG CGG | Dox | (8,6) | Fit failed | Fit failed |
| CCG CGG | Dox | (8,7) | 2.179 | 1.345 |
| CCG CGG | Dox | (9,4) | 9.128 | 0.565 |
| CCG CGG | Dox | (9,5+10,3) | Fit failed | 1.323 |
| CCG CGG | Epi | (10,2) | Fit failed | Fit failed |
| CCG CGG | Epi | (7,5) | 6.334 | Fit failed |
| CCG CGG | Epi | (7,6) | 2.96 | 0.465 |
| CCG CGG | Epi | (8,6) | Fit failed | Fit failed |
| CCG CGG | Epi | (8,7) | Fit failed | 1.042 |
| CCG CGG | Epi | (9,4) | 3.605 | 0.304 |
| CCG CGG | Epi | (9,5+10,3) | Fit failed | 1.048 |
| CCG CGG | Ida | (10,2) | Fit failed | Fit failed |
| CCG CGG | Ida | (7,5) | 14.59 | Fit failed |

| | | | | |
|---|---|---|---|---|
| CCG CGG | Ida | (7,6) | 3.41 | Fit failed |
| CCG CGG | Ida | (8,6) | 3.799 | Fit failed |
| CCG CGG | Ida | (8,7) | 0.616 | Fit failed |
| CCG CGG | Ida | (9,4) | 5.148 | Fit failed |
| CCG CGG | Ida | (9,5+10,3) | Fit failed | Fit failed |
| $(ATTT)_4$ | Dauno | (10,2) | 7.575 | Fit failed |
| $(ATTT)_4$ | Dauno | (7,5) | 8.831 | 2.214 |
| $(ATTT)_4$ | Dauno | (7,6) | 5.382 | 1.63 |
| $(ATTT)_4$ | Dauno | (8,6) | Fit failed | Fit failed |
| $(ATTT)_4$ | Dauno | (8,7) | Fit failed | Fit failed |
| $(ATTT)_4$ | Dauno | (9,4) | 4.91 | 1.586 |
| $(ATTT)_4$ | Dauno | (9,5+10,3) | 2.635 | Fit failed |
| $(ATTT)_4$ | Dox | (10,2) | 8.646 | 0.134 |
| $(ATTT)_4$ | Dox | (7,5) | 20.49 | 3.393 |
| $(ATTT)_4$ | Dox | (7,6) | Fit failed | 1.589 |
| $(ATTT)_4$ | Dox | (8,6) | Fit failed | Fit failed |
| $(ATTT)_4$ | Dox | (8,7) | Fit failed | Fit failed |
| $(ATTT)_4$ | Dox | (9,4) | Fit failed | 1.523 |
| $(ATTT)_4$ | Dox | (9,5+10,3) | Fit failed | 1.582 |
| $(ATTT)_4$ | Epi | (10,2) | 1.948 | Fit failed |
| $(ATTT)_4$ | Epi | (7,5) | Fit failed | Fit failed |
| $(ATTT)_4$ | Epi | (7,6) | Fit failed | 0.967 |
| $(ATTT)_4$ | Epi | (8,6) | Fit failed | Fit failed |
| $(ATTT)_4$ | Epi | (8,7) | Fit failed | Fit failed |
| $(ATTT)_4$ | Epi | (9,4) | Fit failed | 0.965 |

| | | | | |
|---|---|---|---|---|
| $(ATTT)_4$ | Epi | (9,5+10,3) | 1.544 | Fit failed |
| $(ATTT)_4$ | Ida | (10,2) | 9.144 | Fit failed |
| $(ATTT)_4$ | Ida | (7,5) | 11.95 | 3.628 |
| $(ATTT)_4$ | Ida | (7,6) | 7.153 | 1.45 |
| $(ATTT)_4$ | Ida | (8,6) | Fit failed | Fit failed |
| $(ATTT)_4$ | Ida | (8,7) | Fit failed | Fit failed |
| $(ATTT)_4$ | Ida | (9,4) | 10.45 | Fit failed |
| $(ATTT)_4$ | Ida | (9,5+10,3) | 1.648 | Fit failed |
| GGA CGC | Dauno | (10,2) | 20.14 | 1.611 |
| GGA CGC | Dauno | (7,5) | 22.88 | 2.631 |
| GGA CGC | Dauno | (7,6) | 5.576 | 1.517 |
| GGA CGC | Dauno | (8,6) | Fit failed | Fit failed |
| GGA CGC | Dauno | (8,7) | Fit failed | 0.181 |
| GGA CGC | Dauno | (9,4) | 13.14 | 2.274 |
| GGA CGC | Dauno | (9,5+10,3) | 13.37 | Fit failed |
| GGA CGC | Dox | (10,2) | Fit failed | 3.474 |
| GGA CGC | Dox | (7,5) | 13.72 | 4.133 |
| GGA CGC | Dox | (7,6) | 3.527 | 2.145 |
| GGA CGC | Dox | (8,6) | Fit failed | Fit failed |
| GGA CGC | Dox | (8,7) | 15.15 | 0.691 |
| GGA CGC | Dox | (9,4) | 6.171 | 2.202 |
| GGA CGC | Dox | (9,5+10,3) | 42.21 | 4.198 |
| GGA CGC | Epi | (10,2) | 37.84 | 1.724 |
| GGA CGC | Epi | (7,5) | Fit failed | 2.715 |
| GGA CGC | Epi | (7,6) | Fit failed | 1.836 |
| GGA CGC | Epi | (8,6) | Fit failed | Fit failed |
| GGA CGC | Epi | (8,7) | 1.586 | 0.687 |
| GGA CGC | Epi | (9,4) | 2.634 | 1.379 |
| GGA CGC | Epi | (9,5+10,3) | 18.51 | 0.712 |
| GGA CGC | Ida | (10,2) | 69.76 | 1.617 |
| GGA CGC | Ida | (7,5) | 29.28 | 3.251 |

| | | | | |
|---|---|---|---|---|
| GGA CGC | Ida | (7,6) | 6.121 | 1.531 |
| GGA CGC | Ida | (8,6) | Fit failed | Fit failed |
| GGA CGC | Ida | (8,7) | 1.69 | Fit failed |
| GGA CGC | Ida | (9,4) | 4.328 | 1.107 |
| GGA CGC | Ida | (9,5+10,3) | 3.888 | 0.791 |
| $(TAT)_6$ | Dauno | (10,2) | 2.221 | Fit failed |
| $(TAT)_6$ | Dauno | (7,5) | 2.417 | Fit failed |
| $(TAT)_6$ | Dauno | (7,6) | 3.206 | Fit failed |
| $(TAT)_6$ | Dauno | (8,6) | 3.594 | Fit failed |
| $(TAT)_6$ | Dauno | (8,7) | 1.699 | Fit failed |
| $(TAT)_6$ | Dauno | (9,4) | 1.7 | Fit failed |
| $(TAT)_6$ | Dauno | (9,5+10,3) | Fit failed | 3.92 |
| $(TAT)_6$ | Dox | (10,2) | 5.731 | Fit failed |
| $(TAT)_6$ | Dox | (7,5) | 4.86 | 0.307 |
| $(TAT)_6$ | Dox | (7,6) | 5.594 | 0.734 |
| $(TAT)_6$ | Dox | (8,6) | 4.463 | Fit failed |
| $(TAT)_6$ | Dox | (8,7) | 2.369 | Fit failed |
| $(TAT)_6$ | Dox | (9,4) | 3.922 | Fit failed |
| $(TAT)_6$ | Dox | (9,5+10,3) | Fit failed | Fit failed |
| $(TAT)_6$ | Epi | (10,2) | 3.088 | Fit failed |
| $(TAT)_6$ | Epi | (7,5) | 2.751 | Fit failed |
| $(TAT)_6$ | Epi | (7,6) | 4.219 | Fit failed |
| $(TAT)_6$ | Epi | (8,6) | 3.165 | Fit failed |
| $(TAT)_6$ | Epi | (8,7) | 1.956 | Fit failed |
| $(TAT)_6$ | Epi | (9,4) | 1.969 | Fit failed |

| | | | | |
|---|---|---|---|---|
| $(TAT)_6$ | Epi | (9,5+10,3) | Fit failed | Fit failed |
| $(TAT)_6$ | Ida | (10,2) | 9.941 | Fit failed |
| $(TAT)_6$ | Ida | (7,5) | 5.397 | Fit failed |
| $(TAT)_6$ | Ida | (7,6) | 5.196 | Fit failed |
| $(TAT)_6$ | Ida | (8,6) | 4.417 | Fit failed |
| $(TAT)_6$ | Ida | (8,7) | Fit failed | Fit failed |
| $(TAT)_6$ | Ida | (9,4) | 3.738 | Fit failed |
| $(TAT)_6$ | Ida | (9,5+10,3) | Fit failed | 4.813 |
| $(GTT)_3G$ | Dauno | (10,2) | 4.39 | Fit failed |
| $(GTT)_3G$ | Dauno | (7,5) | 5.759 | Fit failed |
| $(GTT)_3G$ | Dauno | (7,6) | 3.72 | Fit failed |
| $(GTT)_3G$ | Dauno | (8,6) | 3.895 | Fit failed |
| $(GTT)_3G$ | Dauno | (8,7) | Fit failed | Fit failed |
| $(GTT)_3G$ | Dauno | (9,4) | 4.065 | 27.99 |
| $(GTT)_3G$ | Dauno | (9,5+10,3) | Fit failed | 22.58 |
| $(GTT)_3G$ | Dox | (10,2) | 4.682 | Fit failed |
| $(GTT)_3G$ | Dox | (7,5) | 4.913 | Fit failed |
| $(GTT)_3G$ | Dox | (7,6) | 5.042 | Fit failed |
| $(GTT)_3G$ | Dox | (8,6) | 4.234 | Fit failed |
| $(GTT)_3G$ | Dox | (8,7) | Fit failed | Fit failed |
| $(GTT)_3G$ | Dox | (9,4) | 5.03 | Fit failed |
| $(GTT)_3G$ | Dox | (9,5+10,3) | Fit failed | Fit failed |
| $(GTT)_3G$ | Epi | (10,2) | 2.175 | Fit failed |
| $(GTT)_3G$ | Epi | (7,5) | 1.559 | Fit failed |

| | | | | |
|---|---|---|---|---|
| $(GTT)_3G$ | Epi | (7,6) | Fit failed | Fit failed |
| $(GTT)_3G$ | Epi | (8,6) | 1.83 | Fit failed |
| $(GTT)_3G$ | Epi | (8,7) | Fit failed | Fit failed |
| $(GTT)_3G$ | Epi | (9,4) | 2.831 | Fit failed |
| $(GTT)_3G$ | Epi | (9,5+10,3) | Fit failed | Fit failed |
| $(GTT)_3G$ | Ida | (10,2) | 3.075 | Fit failed |
| $(GTT)_3G$ | Ida | (7,5) | 3.547 | Fit failed |
| $(GTT)_3G$ | Ida | (7,6) | 2.542 | Fit failed |
| $(GTT)_3G$ | Ida | (8,6) | 2.708 | Fit failed |
| $(GTT)_3G$ | Ida | (8,7) | Fit failed | Fit failed |
| $(GTT)_3G$ | Ida | (9,4) | 3.109 | Fit failed |
| $(GTT)_3G$ | Ida | (9,5+10,3) | 6.145 | Fit failed |
| $(GT)_6$ | Dauno | (10,2) | 11.09 | 0.452 |
| $(GT)_6$ | Dauno | (7,5) | 6.947 | Fit failed |
| $(GT)_6$ | Dauno | (7,6) | 5.26 | Fit failed |
| $(GT)_6$ | Dauno | (8,6) | 9.435 | Fit failed |
| $(GT)_6$ | Dauno | (8,7) | 2.386 | 40.04 |
| $(GT)_6$ | Dauno | (9,4) | 4.707 | Fit failed |
| $(GT)_6$ | Dauno | (9,5+10,3) | Fit failed | Fit failed |
| $(GT)_6$ | Dox | (10,2) | Fit failed | 0.958 |
| $(GT)_6$ | Dox | (7,5) | 8.944 | 0.761 |
| $(GT)_6$ | Dox | (7,6) | 3.967 | 0.752 |
| $(GT)_6$ | Dox | (8,6) | 14.6 | Fit failed |
| $(GT)_6$ | Dox | (8,7) | Fit failed | Fit failed |
| $(GT)_6$ | Dox | (9,4) | 4.934 | 0.396 |
| $(GT)_6$ | Dox | (9,5+10,3) | 3.293 | 1.954 |

| | | | | |
|---|---|---|---|---|
| $(GT)_6$ | Epi | (10,2) | Fit failed | Fit failed |
| $(GT)_6$ | Epi | (7,5) | 3.296 | Fit failed |
| $(GT)_6$ | Epi | (7,6) | Fit failed | Fit failed |
| $(GT)_6$ | Epi | (8,6) | 2.806 | Fit failed |
| $(GT)_6$ | Epi | (8,7) | Fit failed | Fit failed |
| $(GT)_6$ | Epi | (9,4) | 2.036 | Fit failed |
| $(GT)_6$ | Epi | (9,5+10,3) | 2.838 | Fit failed |
| $(GT)_6$ | Ida | (10,2) | 74.69 | Fit failed |
| $(GT)_6$ | Ida | (7,5) | Fit failed | Fit failed |
| $(GT)_6$ | Ida | (7,6) | 4.629 | Fit failed |
| $(GT)_6$ | Ida | (8,6) | 9.828 | Fit failed |
| $(GT)_6$ | Ida | (8,7) | 3.317 | Fit failed |
| $(GT)_6$ | Ida | (9,4) | 4.741 | Fit failed |
| $(GT)_6$ | Ida | (9,5+10,3) | Fit failed | Fit failed |
| $(CTG)_3C$ | Dauno | (10,2) | 53.73 | 0.426 |
| $(CTG)_3C$ | Dauno | (7,5) | 60.34 | 0.809 |
| $(CTG)_3C$ | Dauno | (7,6) | 39.24 | 0.207 |
| $(CTG)_3C$ | Dauno | (8,6) | 37.25 | Fit failed |
| $(CTG)_3C$ | Dauno | (8,7) | Fit failed | Fit failed |
| $(CTG)_3C$ | Dauno | (9,4) | 31.47 | Fit failed |
| $(CTG)_3C$ | Dauno | (9,5+10,3) | 2.571 | Fit failed |
| $(CTG)_3C$ | Dox | (10,2) | 64.13 | 1.198 |
| $(CTG)_3C$ | Dox | (7,5) | 40.21 | 2.743 |
| $(CTG)_3C$ | Dox | (7,6) | 43.88 | 0.833 |
| $(CTG)_3C$ | Dox | (8,6) | 25.2 | 0.62 |
| $(CTG)_3C$ | Dox | (8,7) | Fit failed | 2.154 |
| $(CTG)_3C$ | Dox | (9,4) | 49.21 | 0.634 |

| | | | | |
|---|---|---|---|---|
| $(CTG)_3C$ | Dox | (9,5+10,3) | 2.576 | 1.104 |
| $(CTG)_3C$ | Epi | (10,2) | 11.41 | 0.633 |
| $(CTG)_3C$ | Epi | (7,5) | 7.849 | 3.458 |
| $(CTG)_3C$ | Epi | (7,6) | 14.43 | 0.693 |
| $(CTG)_3C$ | Epi | (8,6) | 13.16 | 0.318 |
| $(CTG)_3C$ | Epi | (8,7) | Fit failed | 1.829 |
| $(CTG)_3C$ | Epi | (9,4) | 7.53 | 0.262 |
| $(CTG)_3C$ | Epi | (9,5+10,3) | Fit failed | 1.114 |
| $(CTG)_3C$ | Ida | (10,2) | 27.58 | Fit failed |
| $(CTG)_3C$ | Ida | (7,5) | 50.46 | Fit failed |
| $(CTG)_3C$ | Ida | (7,6) | 22.63 | Fit failed |
| $(CTG)_3C$ | Ida | (8,6) | 27.6 | Fit failed |
| $(CTG)_3C$ | Ida | (8,7) | Fit failed | Fit failed |
| $(CTG)_3C$ | Ida | (9,4) | 10.35 | Fit failed |
| $(CTG)_3C$ | Ida | (9,5+10,3) | 12.55 | Fit failed |
| $(T)_{12}$ | Dauno | (10,2) | 2.577 | Fit failed |
| $(T)_{12}$ | Dauno | (7,5) | 1.928 | Fit failed |
| $(T)_{12}$ | Dauno | (7,6) | 2.27 | Fit failed |
| $(T)_{12}$ | Dauno | (8,6) | 1.847 | 5.854 |
| $(T)_{12}$ | Dauno | (8,7) | 1.138 | Fit failed |
| $(T)_{12}$ | Dauno | (9,4) | 1.975 | Fit failed |
| $(T)_{12}$ | Dauno | (9,5+10,3) | 0.302 | Fit failed |
| $(T)_{12}$ | Dox | (10,2) | 3.714 | Fit failed |
| $(T)_{12}$ | Dox | (7,5) | 3.982 | Fit failed |
| $(T)_{12}$ | Dox | (7,6) | 2.971 | Fit failed |
| $(T)_{12}$ | Dox | (8,6) | 2.473 | Fit failed |

| | | | | |
|---|---|---|---|---|
| $(T)_{12}$ | Dox | (8,7) | 1.596 | Fit failed |
| $(T)_{12}$ | Dox | (9,4) | 3.576 | 12.16 |
| $(T)_{12}$ | Dox | (9,5+10,3) | Fit failed | Fit failed |
| $(T)_{12}$ | Epi | (10,2) | 1.657 | Fit failed |
| $(T)_{12}$ | Epi | (7,5) | 0.999 | Fit failed |
| $(T)_{12}$ | Epi | (7,6) | 1.434 | Fit failed |
| $(T)_{12}$ | Epi | (8,6) | 1.197 | Fit failed |
| $(T)_{12}$ | Epi | (8,7) | 0.856 | 4.518 |
| $(T)_{12}$ | Epi | (9,4) | 1.259 | Fit failed |
| $(T)_{12}$ | Epi | (9,5+10,3) | Fit failed | Fit failed |
| $(T)_{12}$ | Ida | (10,2) | 3.097 | Fit failed |
| $(T)_{12}$ | Ida | (7,5) | 1.944 | Fit failed |
| $(T)_{12}$ | Ida | (7,6) | 2.351 | Fit failed |
| $(T)_{12}$ | Ida | (8,6) | 2.848 | Fit failed |
| $(T)_{12}$ | Ida | (8,7) | 1.49 | Fit failed |
| $(T)_{12}$ | Ida | (9,4) | 2.571 | Fit failed |
| $(T)_{12}$ | Ida | (9,5+10,3) | Fit failed | Fit failed |
| $(GT)_{30}$ | Dauno | (10,2) | 8.105 | 0.217 |
| $(GT)_{30}$ | Dauno | (7,5) | 9.52 | 0.291 |
| $(GT)_{30}$ | Dauno | (7,6) | 9.743 | 0.226 |
| $(GT)_{30}$ | Dauno | (8,6) | 5.318 | Fit failed |
| $(GT)_{30}$ | Dauno | (8,7) | 3.174 | Fit failed |
| $(GT)_{30}$ | Dauno | (9,4) | 4.968 | Fit failed |
| $(GT)_{30}$ | Dauno | (9,5+10,3) | Fit failed | Fit failed |
| $(GT)_{30}$ | Dox | (10,2) | Fit failed | 0.581 |
| $(GT)_{30}$ | Dox | (7,5) | 11.07 | 1.34 |

| | | | | |
|---|---|---|---|---|
| $(GT)_{30}$ | Dox | (7,6) | Fit failed | 0.886 |
| $(GT)_{30}$ | Dox | (8,6) | 9.915 | 0.285 |
| $(GT)_{30}$ | Dox | (8,7) | 4.676 | 2.959 |
| $(GT)_{30}$ | Dox | (9,4) | 3.331 | 0.239 |
| $(GT)_{30}$ | Dox | (9,5+10,3) | 3.69 | 1.535 |
| $(GT)_{30}$ | Epi | (10,2) | Fit failed | Fit failed |
| $(GT)_{30}$ | Epi | (7,5) | 2.066 | Fit failed |
| $(GT)_{30}$ | Epi | (7,6) | Fit failed | Fit failed |
| $(GT)_{30}$ | Epi | (8,6) | 3.532 | Fit failed |
| $(GT)_{30}$ | Epi | (8,7) | 2.776 | Fit failed |
| $(GT)_{30}$ | Epi | (9,4) | 1.216 | Fit failed |
| $(GT)_{30}$ | Epi | (9,5+10,3) | 42.5 | Fit failed |
| $(GT)_{30}$ | Ida | (10,2) | 8.464 | Fit failed |
| $(GT)_{30}$ | Ida | (7,5) | 10.49 | Fit failed |
| $(GT)_{30}$ | Ida | (7,6) | 9.884 | Fit failed |
| $(GT)_{30}$ | Ida | (8,6) | 3.695 | Fit failed |
| $(GT)_{30}$ | Ida | (8,7) | 3.307 | Fit failed |
| $(GT)_{30}$ | Ida | (9,4) | 5.324 | Fit failed |
| $(GT)_{30}$ | Ida | (9,5+10,3) | Fit failed | Fit failed |

**Supplementary Table 3.** Performance of SVM models for binary concentration classification in buffer and biological matrices in PCA plot. (CV: cross-validation for training; Validation: synthetic urine and sweat)

| | **CV accuracy** | **Test accuracy** | **Validation accuracy** |
|---|---|---|---|
| **Daunorubicin** | 1.0000 ± 0.0000 | 1.0000 | 1.0000 |
| **Doxorubicin** | 1.0000 ± 0.0000 | 0.5000 | 0.5000 |
| **Epirubicin** | 1.0000 ± 0.0000 | 0.5000 | 0.5000 |
| **Idarubicin** | 1.0000 ± 0.0000 | 1.0000 | 1.0000 |

**Supplementary Table 4.** Top 10 ssDNA-(n,m) combinations with highest absolute PC1 loadings for each anthracycline.

| | Feature | PC1 | PC2 | PC3 |
|---|---|---|---|---|
| **Daunorubicin** | $(TAT)_6$*(10,2)_*wl* | 0.851983 | -0.07454 | -0.17132 |
| | $(TAT)_6$*(9,4)_*wl* | 0.382835 | -0.06947 | 0.53657 |
| | $(TAT)_6$*(7,5)_*wl* | 0.143268 | 0.003955 | 0.051422 |
| | $(TAT)_6$*(9,4)_*int* | 0.11432 | 0.488973 | -0.24496 |
| | $(T)_{12}$*(9,4)_*wl* | 0.112942 | -0.15777 | -0.52361 |
| | $(GT)_{15}$*(8,7)_*wl* | 0.106332 | -0.08085 | -0.0621 |
| | GGACGC*(7,6)_*int* | -0.09681 | -0.07074 | 0.010415 |
| | TTATATTATATT*(9,5)+(10,3)_*wl* | 0.094292 | -0.07513 | -0.18019 |
| | TTATATTATATT*(7,6)_*int* | 0.067505 | 0.314878 | -0.09441 |
| | $(GT)_{15}$*(8,6)_*wl* | 0.058032 | 0.034321 | -0.00064 |
| **Doxorubicin** | $(T)_{12}$*(9,4)_*int* | 0.901616 | -0.13951 | 0.067469 |
| | $(T)_{12}$*(7,6)_*int* | 0.30983 | -0.08299 | 0.063758 |
| | $(T)_{12}$*(7,5)_*int* | 0.172457 | -0.07544 | 0.045074 |
| | CCGCGG*(9,5)+(10,3)_*int* | -0.07134 | -0.24448 | 0.294141 |
| | $(T)_{12}$*(7,5)_*wl* | 0.069365 | 0.102523 | -0.15449 |

| | | | | |
|---|---|---|---|---|
| | $(GTT)_3G$*(7,5)_*wl* | 0.066429 | 0.126433 | -0.0506 |
| | $(TCG)_4TC$*(9,5)+(10,3)_*wl* | 0.053682 | 0.063033 | 0.027989 |
| | $(TCG)_4TC$*(7,6)_*wl* | -0.04795 | -0.17822 | 0.223415 |
| | GGACGC*(7,6)_*int* | -0.0443 | -0.23969 | 0.153528 |
| | $(T)_{12}$*(7,6)_*wl* | 0.042971 | 0.015118 | -0.06474 |
| **Epirubicin** | $(TAT)_6$*(7,6)_*int* | 0.408366 | 0.090613 | -0.07485 |
| | $(GTT)_3G$*(9,5)+(10,3)_*int* | 0.380723 | 0.080929 | -0.10661 |
| | $(TAT)_6$*(9,4)_*int* | 0.309901 | 0.049348 | -0.05938 |
| | TTATATTATATT*(7,5)_*int* | 0.250937 | 0.09675 | 0.046434 |
| | $(GTT)_3G$*(7,6)_*int* | 0.196542 | 0.018519 | -0.01875 |
| | TTATATTATATT*(9,4)_*int* | 0.193264 | 0.094664 | 0.101211 |
| | $(TCG)_4TC$*(9,4)_*int* | 0.187575 | 0.098859 | -0.13959 |
| | $(GTT)_3G$*(9,4)_*int* | 0.162952 | 0.002705 | 0.007055 |
| | $(GT)_{15}$*(7,6)_*int* | 0.15612 | 0.004774 | -0.05593 |
| | $(GT)_{15}$*(9,4)_*int* | 0.149052 | 0.002878 | -0.04324 |
| **Idarubicin** | $(GT)_{30}$*(9,4)_*int* | 0.315611 | 0.068914 | -0.17637 |
| | CCGCGG*(9,5)+(10,3)_*int* | 0.299782 | 0.130549 | -0.16999 |

| | | | | |
|---|---|---|---|---|
| | $(GT)_{30}$*(7,6)_*int* | 0.282599 | 0.162562 | -0.18965 |
| | $(GTT)_3$G*(7,6)_*int* | 0.265926 | 0.02377 | -0.14202 |
| | $(GTT)_3$G*(9,4)_*int* | 0.251676 | 0.00191 | -0.16277 |
| | $(GT)_{30}$*(7,5)_*int* | 0.208016 | 0.135192 | -0.10672 |
| | $(GT)_6$*(7,5)_*int* | 0.16618 | -0.0013 | -0.02796 |
| | $(GT)_{30}$*(10,2)_*int* | 0.15978 | 0.109552 | -0.0733 |
| | CCGCGG*(7,6)_*int* | 0.151966 | 0.119974 | -0.03878 |
| | $(GT)_{15}$*(7,6)_*int* | 0.151453 | 0.107696 | -0.02382 |
